# Time-, tissue- and treatment-associated heterogeneity in tumour-residing migratory DCs

**DOI:** 10.1101/2023.07.03.547454

**Authors:** Colin YC Lee, Bethany C Kennedy, Nathan Richoz, Isaac Dean, Zewen K Tuong, Fabrina Gaspal, Zhi Li, Claire Willis, Tetsuo Hasegawa, Sarah K Whiteside, David A Posner, Gianluca Carlesso, Scott A Hammond, Simon J Dovedi, Rahul Roychoudhuri, David R Withers, Menna R Clatworthy

## Abstract

Tumour dendritic cells (DCs) internalise antigen and upregulate CCR7, which directs their migration to tumour-draining lymph nodes (dLN). CCR7 expression is coupled to a maturation programme enriched in regulatory molecule expression, including PD-L1, termed mRegDC. However, the spatio- temporal dynamics and role of mRegDCs in anti-tumour immune responses remain unclear. Using photoconvertible mice to precisely track DC migration, we found that mRegDCs were the dominant DC population arriving in the dLN, but a subset remained tumour-resident despite CCR7 expression. These tumour-retained mRegDCs were phenotypically and transcriptionally distinct from their dLN counterparts and were heterogeneous. Specifically, they demonstrated a progressive reduction in the expression of antigen presentation and pro-inflammatory transcripts with more prolonged tumour dwell-time. Tumour mRegDCs spatially co-localised with PD-1^+^CD8^+^ T cells in human and murine solid tumours. Following anti-PD-L1 treatment, tumour-residing mRegDCs adopted a state enriched in lymphocyte stimulatory molecules, including OX40L, which was capable of augmenting anti- tumour cytolytic activity. Altogether, these data uncover previously unappreciated heterogeneity in mRegDCs that may underpin a variable capacity to support intratumoural cytotoxic T cells, and provide insights into their role in cancer immunotherapy.

## Introduction

Dendritic cells (DCs) capture tumour antigens and upregulate the chemokine receptor CCR7, which directs their migration to secondary lymphoid organs where they present antigens to T cells^1–3^. Two major conventional DC (cDC) subsets have been identified in tumours; cDC1 that specialise in cross- presenting tumour antigens to CD8^+^ T cells and are associated with improved survival^4^, and cDC2 that present exogenous antigens to CD4^+^ T cells and have variable associations with cancer prognosis and treatment responses^1^. The recent application of single-cell technologies to tissue samples has enabled the distinction of an intratumoral DC population that may arise from both cDC1 and cDC2, but demonstrates a conserved phenotype and transcriptional programme characterised by the expression of LAMP3, genes consistent with a mature and migratory state (CCR7, CD40, IL12), and immunoregulatory molecules including PD-L1 and PD-L2^5^. This DC subset or state has been variously labelled as “migratory DCs”, “mature DCs enriched in immunoregulatory molecules” (mRegDCs, the term we adopted in the current study), as well as “LAMP3^+^ DCs”, “mature DCs”, or “cDC3”^6–8^. Regardless of nomenclature, single-cell atlasing efforts have identified mRegDCs in multiple human cancers^9–11^, and the acquisition of this maturation programme appears dependent on the uptake of tumour antigens^5^.

Intriguingly, despite the conserved expression of co-inhibitory molecules such as PD-L1 in mRegDCs, the expression of the mRegDC-associated marker LAMP3 is associated with improved prognosis in breast, lung cancer and metastatic melanoma^12–15^. However, the precise contribution of mRegDCs to anti-tumour responses, and whether they have a positive or negative effect on disease outcomes remains unclear. Certainly, mRegDCs are likely to be key targets of immune checkpoint blockade (ICB) by virtue of their high expression of co-inhibitory molecules. Consistent with this, murine studies demonstrate the importance of DC-expressed PD-L1 in determining anti-tumour cytotoxic responses to subcutaneously tumours^16,17^, and the application of anti-IL4 to increase mRegDCs enhanced responses to anti-PD-L1 in a non-small cell lung cancer (NSCLC) model^5^.

mRegDCs express high levels of CCR7, enabling trafficking from the tumour to draining lymph nodes (dLN)^2,3,5^. Indeed, LAMP3^+^ DCs have been identified in tumour-dLN, where they may activate tumour antigen-specific T cells^18^. However, LAMP3-expressing DCs have also been described within tertiary lymphoid structures (TLS) in tumours^12,15^, suggesting that some mature DCs may be retained in tumours for prolonged periods. Thus, the precise temporal dynamic behaviour of mRegDCs, the extent to which they act within the tumour versus dLN, and how these dynamics and site-specific cellular interactions are influenced by ICB remain to be clarified. The concept that prolonged residence within tumours might influence mRegDC fate and function is worthy of consideration given the known effects of the tumour microenvironment (TME) on other immune cell populations. For example, CD8^+^ T cells transition to a so-called ‘exhausted’ state with prolonged dwell-time in the tumour^19^, with three defining characteristics; reduced effector function, sustained expression of inhibitory molecules such as PD-1, and a transcriptional state distinct from that of functional effector cells^20^. These features have also been observed in other tumour-resident immune cells such as natural killer (NK) cells^21,22^.

Here, we used a photo-tracking mouse model, combined with single-cell RNA sequencing (scRNA- seq), confocal imaging and spatial transcriptomics, in mouse and human tissues, to interrogate the spatio-temporal dynamics of mRegDCs, their roles within the tumour, and the effects of ICB. We found mRegDCs were heterogeneous, influenced by duration of tumour residence, location (tumour versus dLN), and anti-PDL1 treatment. Strikingly, tumour-retained mRegDCs showed similar expression of CCR7 to those that migrated the dLN but took on an increasingly “exhausted” transcriptional profile with more prolonged tumour dwell-time. These intratumoural mRegDCs co-localised with cytotoxic CD8^+^ T cells, and anti-PD-L1 treatment enhanced their expression of several important T cell-stimulatory molecules. mRegDC-CD8^+^ T cell engagement and their augmentation by ICB was confirmed across a range of human cancers. Altogether, these data provide further insight to tumour mRegDC dynamics and their role in ICB therapy.

## Results

### mRegDC signatures are associated with improved survival in human cancers

The presence of mature LAMP^+^ DCs within lymphoid follicles has been associated with improved prognosis in NSCLC^12^, but the effect of tumour mRegDCs on prognosis in human cancers more broadly has not been examined. To address this, we analysed the transcriptomes of 4,045 human solid tumours from the cancer genome atlas (TCGA)^23^. Enrichment of an mRegDC signature was associated with improved survival not only in lung cancer, but also in cutaneous melanoma, breast, and colorectal cancer (CRC, **Extended Data Fig. 1A**), all of which harbour CCR7^+^ mRegDCs (**Extended Data Fig. 1B-G**)^24–26^. Further analysis of 1,853 human breast tumours from METABRIC^27^ revealed cancer subtype-specific associations with survival (**Extended Data Fig. 1H**). These data suggest that mRegDCs contribute to anti-tumour responses across a range of human cancers.

### mRegDCs are transcriptionally heterogenous, and some remain within the tumour despite CCR7 expression

To investigate the mechanisms by which mRegDCs promote anti-tumour immunity, we sought to characterise their spatio-temporal dynamics. The mRegDC programme is thought to be driven by acquisition of tumour antigen^5^, and we asked whether establishment of this programme inevitably leads to the trafficking of CCR7^+^ mRegDCs to the dLN, or whether some cells remain in the tumour. To address this, we established multiple syngeneic subcutaneous colorectal tumours (MC38, MC38-Ova, CT26) in a photoconvertible Kaede transgenic model that enables site-specific temporal labelling of cells within the tumour^28^. Tumours were transcutaneously photoconverted on day 13, converting all infiltrated cells in the tumour only from the default green fluorescence (Kaede-green) to a red fluorescent profile (Kaede-red)^19^. 24-72h following photoconversion, tumours were harvested, enabling the distinction of newly-infiltrating Kaede-green cells and Kaede-red cells retained in the tumour from the point of photo-labelling (**Fig. 1A**). To address the effect of ICB on mRegDCs, we administered anti-PD-L1 antibodies in this model (**Extended Data Fig. 2A**), selecting this target because mRegDCs in CRC showed the highest expression of PD-L1 among immune cells (**Extended Data Fig. 1C**).

**Figure 1.**
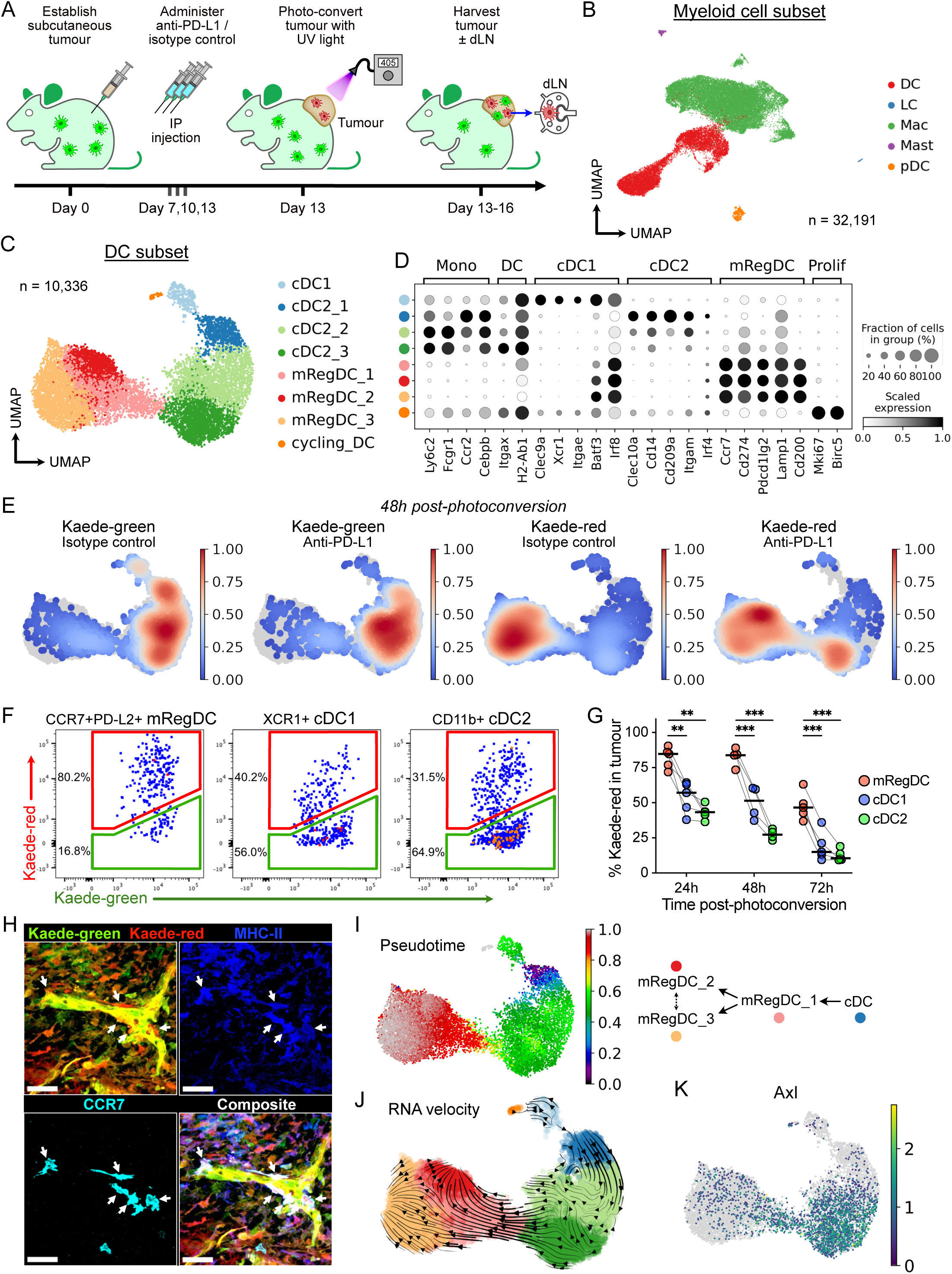
Landscape and temporal dynamics of DCs in murine subcutaneous tumours. (A) Experiment design. Tumours from Kaede transgenic mice were harvested 5h, 24h, 48h, or 72h after photoconversion. (B) UMAP of myeloid cells from scRNA-seq of FACS-sorted CD45^+^TER119^-^ Kaede- green^+^/Kaede-red^+^ cells 48h after photoconversion of subcutaneous MC38-Ova tumours. (C) UMAP of DCs from (B), and canonical marker gene expression of respective clusters (D). (E) Kernel density embedding of DCs by Kaede fluorescence and treatment group. (F) Representative flow cytometry of Kaede fluorescence in DCs subsets from MC38-Ova tumours 48h post-photoconversion. Data is representative of 3 independent experiments (12 independent mice). (G) Flow cytometry of Kaede fluorescence in DCs from MC38-Ova tumours. Points represent independent mice. Paired t-test with FDR correction was used. (H) Representative confocal microscopy images of MC38 tumours 72h after photoconversion. Scale bar, 40μm; arrows, tumour-residing mRegDCs (Kaede- red^+^CCR7^+^MHC-II^+^, dendritic morphology). (I) Tumour DCs ordered by pseudotime, rooted in the cluster with highest proportion of Kaede-green DCs; and proposed pseudotime trajectory. (J) RNA velocity trajectory in tumour DCs. (K) Expression of *Axl*.

scRNA-seq of FACS-isolated Kaede-green^+^ or Kaede-red^+^ immune cells (Kaede^+^CD45^+^Ter119^-^) 48h after photoconversion (**Extended Data Fig. 2B**) generated 80,556 single cell transcriptomes, following rigorous quality control, including 32,191 myeloid cells (**Fig. 1B**, **Extended Data Fig. 2C**). Unbiased clustering of DCs revealed 8 distinct clusters, including cycling DCs, cDC1, cDC2s, and mRegDCs, assigned based on expression of canonical marker genes and similarity to published transcriptomes (**Fig. 1C-D**, **Extended Data Fig. 2D**), with mRegDC showing high expression of *Ccr7*, *Cd274* and *Pdcd1lg2*. Indeed, mRegDCs expressed higher levels of surface PD-L1 than other immune cells (**Extended Data Fig. 2E**).

CCR7^+^PD-L2^+^ mRegDC were a prominent tumour DC population (**Fig. 1C**), and this was confirmed by flow cytometry (**Extended Data Fig. 2F-G**). Surprisingly, tumour mRegDCs were predominantly Kaede-red indicating that they have resided in the tumour for over 48h (**Fig. 1E**, **Extended Data Fig. 2H**). In contrast, cDC1 and cDC2 were mostly Kaede-green, consistent with the conclusion that these newly infiltrating cDCs differentiate into mRegDCs following uptake of tumour antigen^5^. The prevalence of tumour-retained Kaede-red mRegDCs was validated over a time-course in all tumour models, by flow cytometry, where CCR7^+^PD-L2^+^ mRegDCs were the major Kaede-red DC cell-type up to 72h post-photoconversion (**Fig. 1F-G**, **Extended Data Fig. 2I**). Using immunofluorescence (IF) microscopy, we confirmed the presence of Kaede-red MHC-II^+^CCR7^+^ cells (**Fig. 1H**, **Extended Data Fig. 2J-L**).

To explore the relationship between tumour DC subsets, we performed pseudotime analysis rooted in the Kaede-green dominant cluster (cDC2_1). This revealed a trajectory terminating in mRegDC_2/3 that transitioned through an intermediate mRegDC_1 state (**Fig. 1I**, **Extended Data Fig. 3A**), consistent with the Kaede-green/red ratio of clusters that marks tumour dwell in real-time. RNA velocity analysis confirmed the cell-state transition from cDC, through mRegDC_1, to mRegDC_2 or mRegDC_3, which were the endpoints of the velocity trajectory (**Fig 1J**, **Extended Data Fig. 3B**).

To further prove that tumour mRegDC arise from cDC precursors, we examined Kaede-green DCs 5h post-photoconversion, which should harbour few mRegDC because newly-entered DCs would have little time to acquire the mRegDC programme. Indeed, at 5h post-photoconversion the Kaede-green mRegDC:cDC ratio was only 1/100, but this increased to 1/5 72h post-photoconversion (**Extended Data Fig. 3C**). Hence, the mRegDC state emerges over time in the tumour and only after the influx of cDCs. cDC1 or cDC2-defining transcripts were not conserved upon acquisition of the mRegDC programme, for example, *Clec9a* or *Clec10a* expression was lost but *Irf8* and *Batf3* are upregulated ubiquitously (**Fig 1D**). However, retained cDC subset-specific surface marker expression indicated a mixed ontogeny of mRegDCs, with expression of XCR1 (cDC1 origin) and CD11b (cDC2 origin) detectable, and notably, XCR1^+^ mRegDCs were higher among the Kaede-red fraction (**Extended Data Fig. 3D**). These data support the conclusion that both cDC1 and cDC2 maturate to tumour-residing mRegDCs.

Kaede-red^+^ cDC1 and cDC2 showed higher expression of migratory transcripts and *Ciita*, a master regulator of antigen presentation^29^, than Kaede-green^+^ counterparts, suggesting that cDCs mature with duration in the TME (**Extended Data Fig. 3E**). Principal component (PC) analysis showed that Kaede status (green/red) accounted for most of the transcriptional variance in cDC2s (PC1, **Extended Data Fig. 3F**). Genes driving PC1 were enriched in pathways relating to antigen presentation and myeloid differentiation (**Extended Data Fig. 3G**). cDC2_3, the most mature cluster in the cDC2-to-mRegDC trajectory, upregulated *Ciita*, class-II MHC (MHC-II) transcripts, and *Ccr7* (**Extended Data Fig. 3I**). The receptor tyrosine kinase *Axl*, which recognises apoptotic cells and induces the mRegDC programme^5^, was also upregulated as cDC2_1 transitioned to cDC2_3 (**Figure 1K**). Altogether, these data suggest time-associated maturation of cDCs in the TME towards an mRegDC state, and importantly, antigen-charged mRegDCs may reside in the tumour for several days despite the expression of genes involved in tumour egress, including CCR7.

### Migrated mRegDCs are phenotypically distinct from tumour-residing mRegDCs

DCs expressing mRegDC transcripts have been identified in tumour-dLN^5,30^ and CCR7 expression directs migration to the dLN. However, given our demonstration that many mRegDCs acquire prolonged tumour residence, we sought to definitively track tumour DC egress to dLNs. Kaede-red CCR7^+^PD-L2^+^ DCs, which were photo-labelled within the tumour and hence tumour emigrants, were readily detectable in the dLN, but not in the contralateral non-draining LN (ndLN, **Fig. 2A-B**). Indeed, among Kaede-red DCs in the dLN, essentially all were mRegDC, up to 72h post-photoconversion (**Fig. 2C**). scRNA-seq of Kaede-red CD45^+^ cells in tumour-dLNs (integrated with CD45^+^ cells from control LNs) included a prominent *Ccr7^+^Cd274*^+^ mRegDC cluster among myeloid cells (**Fig. 2D**, **Extended Data Fig. 4A**). In this data, 94% of Kaede-red^+^ myeloid cells in the dLN were mRegDCs (**Fig. 2E**). Hence, mRegDC are the dominant myeloid cell tumour emigrants arriving in the dLN.

**Figure 2.**
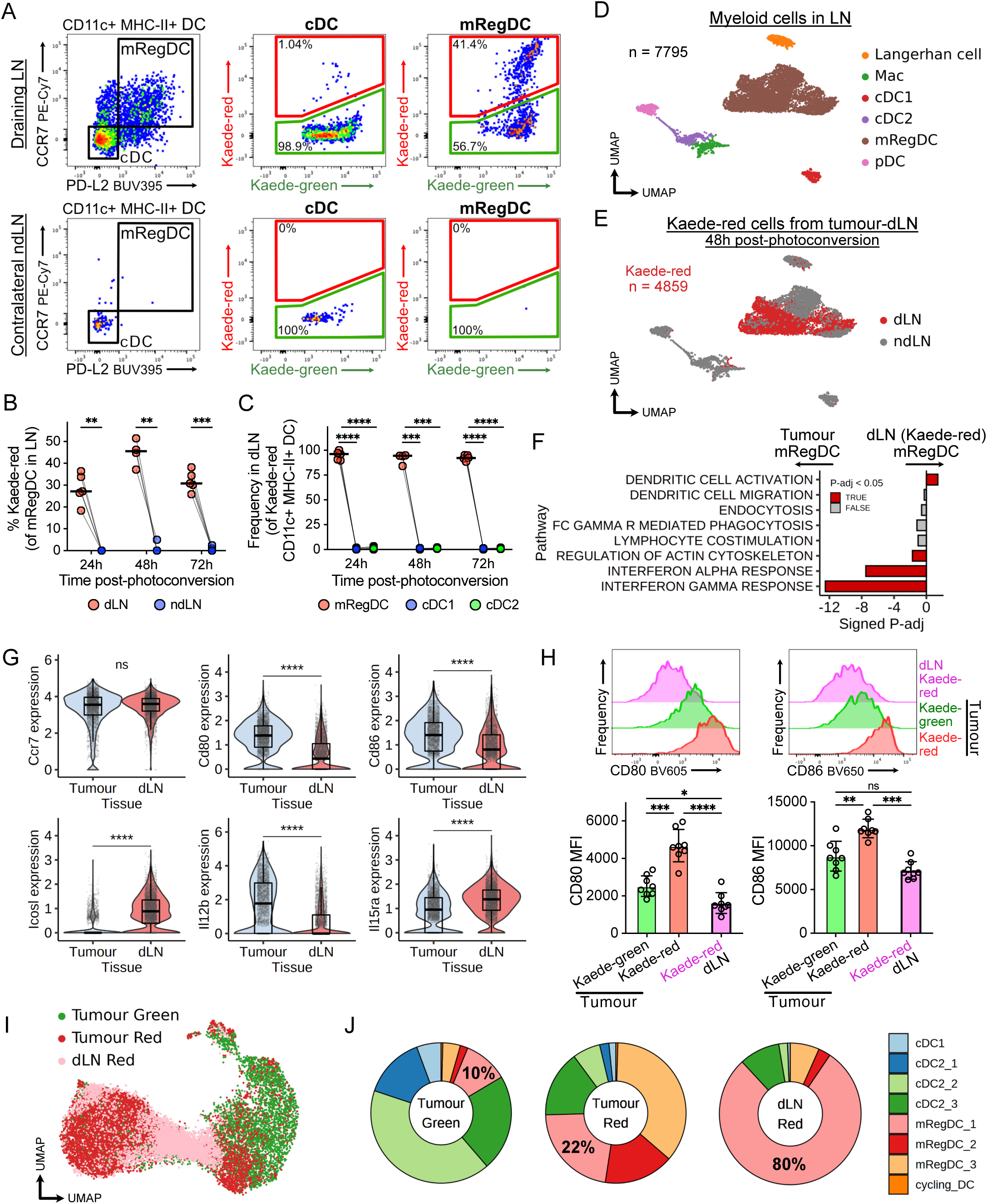
mRegDCs that migrate to the dLN become phenotypically and transcriptionally distinct from tumour-residing populations. (A) Representative flow cytometry of tumour-dLNs and contralateral ndLNs 48h after tumour photoconversion. DCs that originate from photo-flashed tumours carry the Kaede-red fluorescent profile, which enables tracking to LNs. (B) Flow cytometry of Kaede-red cells (tumour emigrants) among mRegDCs in the dLN or ndLN *x* hours after tumour photoconversion. (C) Flow cytometry of Kaede-red DC identity in dLNs. MC38-Ova tumours were used (A- C); points represent independent mice; paired t-test with FDR correction was used (B-C). (D-E) UMAP of myeloid cells from scRNA-seq of FACS-sorted CD45^+^ Kaede-red cells from tumour-dLNs (MC38-Ova) and CD45^+^ cells from control LNs. (F) GSEA of Kaede-red mRegDCs in dLNs versus tumours (isotype control-treated). Signed P-adj indicate log10(adjusted *P*-value) with the direction of enrichment. (G) Expression of selected genes in tumour mRegDCs and Kaede-red mRegDCs in dLNs (isotype control-treated). Wilcoxon rank-sum test was used. (H) Flow cytometry of CD80/86 on mRegDCs from MC38-Ova tumours or dLNs 48h after photoconversion, with representative histograms. Mean MFI ± standard deviation shown; points represent independent mice; paired t-test with FDR correction was used. (I) Integration of scRNA-seq of Kaede-red DCs from the tumour-dLN with tumour DCs and label transfer. (J) Proportion of DCs by tissue and Kaede profile, related to (I).

Compared to Kaede-red tumour emigrants in the dLN, mRegDCs in tumours were enriched in ‘*interferon gamma response*’ genes (**Fig. 2F**). While there was no difference in mRegDC ‘*lymphocyte co-stimulation*’ geneset expression between sites, expression of precise co-stimulatory molecules differed (**Fig. 2F**, **Extended Data Fig. 4B-C**). Specifically, CD80 and CD86 were higher in tumour- residing mRegDCs, confirmed at a protein level, while *Icosl* was significantly higher in migrated mRegDCs in the dLN (**Fig. 2G-H**, **Extended Data Fig. 4B**). Of note, CD80/86 and ICOSL have differing effects on T cell activation, where signalling through CD28 but not ICOS induces IL-2 production to support the clonal expansion of T cells^31^. *Il12b*, which drives anti-tumour cytotoxic T cell activity^32^, was enriched in tumour mRegDCs, while *Il15ra*, associated with DC maturation^33^, was higher in mRegDC in the dLN. Notably, scRNA-seq of B16-F10 tumours and their dLNs^34^ demonstrated the same tissue-associated heterogeneity in mRegDCs (**Extended Data Fig. 4D-G**).

We next asked whether the three transcriptionally distinct tumour mRegDC states contributed equally to the dLN emigrants. To address this, we integrated the single-cell transcriptomes of tumour-originating DCs in the dLN with the tumour DC landscape (**Fig. 2I**). Strikingly, over 80% of Kaede- red DCs from the dLN mapped to the tumour mRegDC_1 cluster (**Fig. 2J**). Only 9% of DCs in the dLN resembled mRegDC_2/3, despite mRegDC_2/3 being the dominant Kaede-red^+^ tumour mRegDC states. Although CCR7 expression remained high in both tumour-residing or migrated mRegDCs, we observed a decrease in transcripts associated with DC chemotaxis and CCR7 signalling in mRegDC_2 and 3 (**Extended Data Fig. 4H-J**). Moreover, CD80 and CD86 expression was upregulated on Kaede- red versus Kaede-green tumour mRegDCs, with expression levels on tumour Kaede-green mRegDCs similar to migrated mRegDCs in matched dLNs (**Fig 2H**). Therefore, successful LN emigrants resemble newly-formed mRegDC, and tumour-retained mRegDCs acquire a distinct phenotype with prolonged tumour dwell-time. Altogether, these data suggest that mRegDC_1, which have most recently acquired the mRegDC programme, are the main population to seed the dLN. Hence, we propose that mRegDC_2 and mRegDC_3 are terminal, tumour-residing mRegDC states, and DCs that have transitioned beyond the intermediate mRegDC_1 state become increasingly unlikely to egress.

### Tumour-retained mRegDCs progress towards an “exhausted”-like state but is attenuated by anti-PD-L1 treatment

Since tumour-retained mRegDCs are reduced in their migratory capacity, we sought to assess how their transcriptional programmes changed with increasing time in the tumour. During the progression from mRegDC_1 to mRegDC_3, there was a decrease in MHC-II expression, including class-II invariant chain *Cd74* (**Fig. 3A**, **Extended Data Fig. 5A**). Concomitantly, DC antigen presentation genes decreased as cells transitioned towards the tumour-retained mRegDC_2/3 states (**Fig. 3B-C**). This included downregulation of molecules involved in antigen processing (*Ctsb*, *Ctsl*, *Ifi30*, *Lgmn*, *Psme2*), transport (*Tap1*, *Tap2*), chaperones (*Hsp1a1*, *Hspa2*, *Hspa8*), MHC loading (*H2-DMa*, *H2-Oa*, *Tapbp*, *Calr*) and *Ciita* (**Extended Data Fig. 5B-C**).

**Figure 3.**
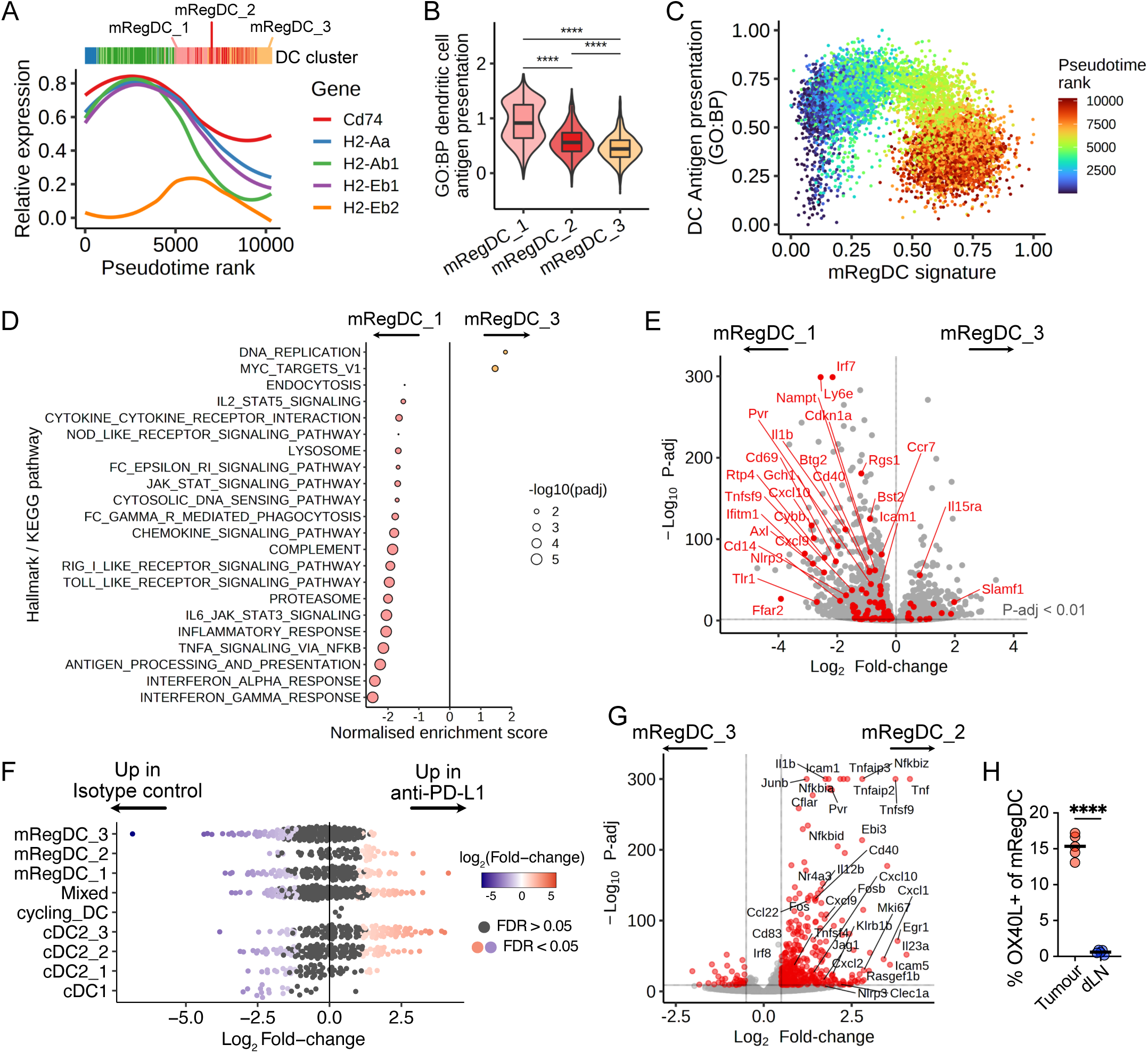
Tumour-residing mRegDCs undertake an “exhausted” state with duration in the tumour, attenuated in anti-PD-L1 treatment. (A) Expression of MHC-II transcripts and *Cd74* over pseudotime in tumour DCs. Local regression (loess) was fit to scaled expression values. (B) Gene signature scores for “*GO:BP dendritic cell antigen processing and presentation (GO:0002468)*” across mRegDC clusters. Wilcoxon rank-sum test was used. (C) Gene signature scores (scaled) of individual cells for “*GO:BP dendritic cell antigen processing and presentation (GO:0002468)*” and mRegDC signature genes, coloured by pseudotime. (D) GSEA of mRegDC_3 versus mRegDC_1. Only significant pathways (*P*-adj < 0.05) shown. (E) Differential gene expression between mRegDC_1 and mRegDC_3. Significant DEGs from “*Hallmark inflammatory response*” are highlighted in red (*P*-adj < 0.01). (F) Milo differential abundance analysis. Bee-swarm plot shows treatment-associated differences in overlapping cellular neighbourhoods (points). Differentially abundant neighbourhoods at FDR < 0.05 are coloured. ‘Mixed’ refers to neighbourhoods where cells do not predominantly (>70%) belong to a single cluster. (G) Differential gene expression between mRegDC_2 and mRegDC_3. DEGs (*P*-adj < 0.01, log_2_Fold- change > 0.5) are coloured. (H) Flow cytometry of OX40L expression on mRegDC from MC38-Ova tumours and their dLN. Paired t- test was used.

Moreover, genes involved in innate immune function and response to inflammatory stimuli, including ‘*interferon gamma/alpha response*’, ‘*inflammatory response*’, ‘*cytokine-cytokine receptor interaction*’ and ‘*Toll-like receptor signalling*’ pathways, significantly decreased as cells transitioned from mRegDC_1 to mRegDC_3 (**Fig. 3D**). Specifically, mRegDC_3 showed reduced expression of several innate immune response genes (*Nlrp3*, *Tlr1*, *Cd14*), genes involved in DC activation and migration (*Axl*, *Ccr7*, *Rgs1*, *Icam1*), lymphocyte co-stimulation (*Cd40*, *Tnfsf9*, *Pvr*) and cytokines which may recruit and activate immune cells (*Cxcl9*, *Il1b*, *Tnf*) compared to mRegDC_1 (**Fig. 3E**, **Extended Data Fig. 5D**). To identify transcription factors (TF) that accompany the tumour-retained mRegDC state, we performed TF regulon analysis, which revealed that *Tcf7* expression and its activity were upregulated in terminal mRegDC states (**Extended Data Fig. 5E**). Altogether, these data suggest that mRegDCs retained in the tumour undergo a transition, acquiring the transcriptional hallmarks of “exhaustion”, as defined in T cells^20^, namely reduced expression of molecules enabling effector function (e.g. antigen presentation), sustained expression of inhibitory molecules (such as PD-L1/L2), and a transcriptional state distinct from that of functional effector cells (i.e. successful dLN emigrants).

Next, we considered the effects of anti-PD-L1 treatment on tumour DCs, since PD-L1 expression by DCs is essential for effective responses to anti-PD-L1 antibodies^16,17^. In cDC2s from anti-PD-L1- treated tumours, genes involved in inflammation and adaptive immunity were upregulated and expression of *Axl* increased (**Extended Data Fig. 3H-J**). Across all mRegDC states, differential gene expression following anti-PD-L1 drove enrichment of similar pathways, including “*interferon gamma response*”, necessary for effective response to ICB^32^, and “*TNFα signalling via NFκB*” (**Extended Data Fig. 6A-B**).

We then asked whether anti-PD-L1 treatment influenced tumour mRegDC heterogeneity. Assessment of differential abundance showed significant enrichment within mRegDC_2 neighbourhoods following anti-PD-L1 treatment, but relative depletion of mRegDC_3 (**Fig. 3F**), consistent with kernel density embeddings (**Fig 1E**). RNA velocity analysis suggested that this was driven by a preferential transition from mRegDC_1 to mRegDC_2 in anti-PD-L1-treated tumours (**Extended Data Fig. 6C**). Strikingly, the transcriptome of mRegDC_2 was highly activated compared to mRegDC_3, with increased expression of many genes involved in immune signalling, including *Cd40*, *Il1b*, *Tnf*, *Nfkbia*, etc. (**Fig. 3G**). This was reflected in the pathway enrichment analysis, which showed multiple genesets relating to immune activation enriched in mRegDC_2 (**Extended Data Fig. 6D**), potentially augmenting their capacity to promote anti-tumour responses.

Importantly, we identified several lymphocyte activation ligands^32,35,36^ that were significantly upregulated on mRegDC_2 versus other tumour mRegDC states and dLN mRegDCs, such as *Il12b*, *Tnfsf4*, *Tnfsf9*, *Cd70* and *Pvr* (**Extended Data Fig. 6E**). These molecules were also preferentially expressed on tumour-residing mRegDCs in B16-F10 tumours (**Extended Data Fig. 4G**). Using OX40L^Hu-CD4^ (*Tnfsf4*)-reporter mice, we confirmed that some tumour mRegDCs express OX40L, but it was not expressed by tumour cDCs or mRegDCs in the dLN (**Fig. 3H**, **Extended Data** Fig 6F-G). The absence of OX40L expression in the dLN suggests that it is only upregulated by tumour-retained CCR7^+^ DCs, and that OX40L^+^ mRegDC do not emigrate to the dLN, or at least rapidly downregulate it upon tumour exit. Altogether, these data underline that a subset of tumour-retained mRegDCs exhibit a specific, “activated” state, which is enriched following anti-PD-L1 treatment.

### mRegDCs interact with CD8^+^ T cells in human tumours

Given that a substantial proportion of mRegDCs in the tumour failed to migrate to dLN, and that tumour mRegDCs may associate with outcomes, we sought to identify which immune cells mRegDCs may communicate with in the tumour. We deconvolved 521 human CRC transcriptomes^23^ and found the highest correlation between mRegDC and CD8^+^ T cell transcripts (**Fig. 4A**), an observation replicated in melanoma, breast, and lung tumour biopsies (**Fig. 4B**). Furthermore, the *CCR7*^+^*CD274*^+^*PDCD1LG2*^+^ DC cluster (consistent with mRegDCs in human CRC^26^) (**Extended Data Fig. 1B-C**), had stronger predicted interactions with effector CD8^+^ T cells than other myeloid cells, including via CD28, CTLA-4 and PD-1 (*PDCD1*) engagement (**Fig. 4C**). Similar observations were confirmed in scRNA-seq of human breast tumours and melanoma (**Extended Data Fig. 1D-G**)^24,25^.

**Figure 4.**
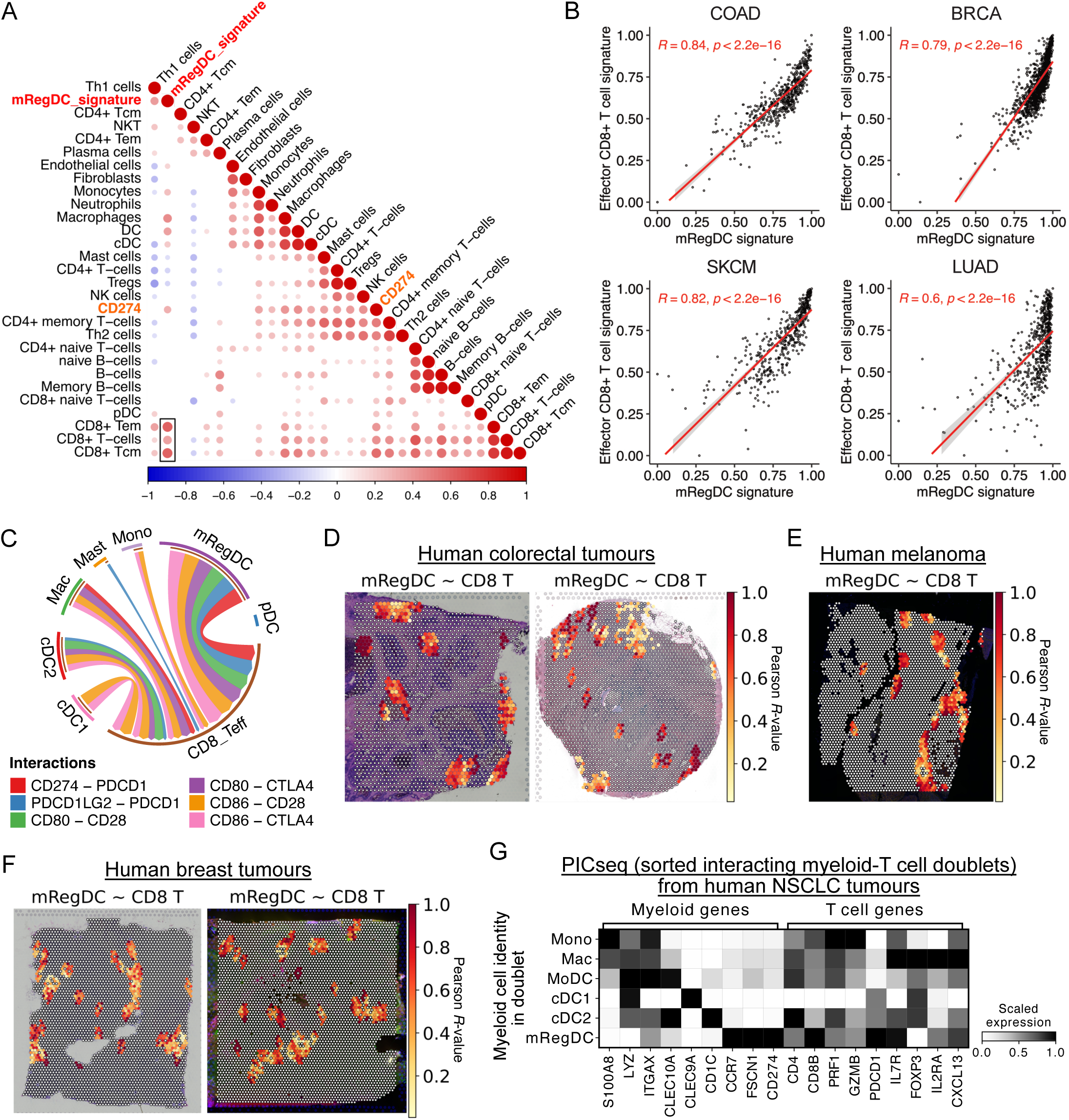
mRegDCs interact with CD8^+^ T cells in human cancers. (A) Pearson correlation between cell proportions, from deconvolution of 521 bulk transcriptomes of colorectal adenocarcinoma biopsies (TCGA), and mRegDC signature genes. Box highlights high correlation between mRegDC and CD8^+^ T cells. Only significant correlations (*p* < 0.05) shown. (B) Pearson correlation between mRegDC signature genes and effector CD8^+^ T cell signature genes in colorectal cancer (COAD), breast cancer (BRCA), cutaneous melanoma (SKCM) and lung adenocarcinoma (LUAD) from TCGA. (C) CellPhoneDB cell-cell communication analysis between myeloid cells and effector CD8^+^ T cells in scRNA-seq of human CRC. Edge width scaled to standardised interaction scores. Only significant interactions (*p* < 0.05) shown. (D-F) Spatial correlation (Pearson R-value) of mRegDC and effector CD8^+^ T cell signature scores in spatial transcriptomics (10X Genomics Visium) of independent human CRC tumour sections (D, *n=* 2), human melanoma section (E, *n=* 1), and independent human breast tumour sections (F, *n=* 2). (G) Expression of selected genes in myeloid-T cell doublets from PICseq of NSCLC tumours, grouped by the myeloid cell identity in each myeloid-T cell doublet.

Effective engagement of cell-surface ligand-receptor pairs require co-localisation of mRegDCs and effector CD8^+^ T cells. Analysis of spatial transcriptomics data from human CRC tumours^37^ revealed hotspots with co-localised expression of mRegDC and effector CD8^+^ T cell genes (**Fig. 4D**, **Extended Data Fig. 7A-B**). Spatial correlation of mRegDC and effector CD8^+^ T cell transcripts was also evident in human melanoma and breast tumours (**Fig. 4E-F**).

Finally, we analysed data from RNA-seq of physically-interacting cells (PICs), consisting of sorted myeloid-T cell doublets from NSCLC^38^. Of note, mRegDC-T cell PICs were more frequent in tumours than normal tissue^38^. We found that mRegDC-CD8^+^ T cell doublets were more frequent than other mRegDC-T cell combinations (**Extended Data Fig. 7C-D**), and PICs containing mRegDC (*CCR7*^+^*FSCN1*^+^*CD274^+^*) highly co-expressed *CD8B, PRF1, GZMB and PDCD1* (**Fig. 4G**), confirming that in NSCLC, mRegDC and effector CD8^+^ T cells physically interact. Altogether, these data suggest that mRegDC-CD8^+^ T cell interaction is conserved across multiple human solid tumours. The prolonged tumour dwell-time of mRegDCs, which maintain high levels of PD-1 ligand expression, but downregulate expression of genes enabling effector function, suggests that these cellular interactions are potentially deleterious. Specifically, tumour-retained mRegDC may regulate the activation and expansion of anti-tumour cytotoxic T cells, but they could also be important targets of cancer immunotherapy.

### Anti-PD-L1 promotes immunogenic mRegDC-CD8^+^ T cell interactions

To investigate the effects of anti-PD-L1 on mRegDC-CD8 T cell interactions in tumours, we first analysed scRNA-seq of TILs (**Extended Data Fig. 8A-B**), focussing on PD-1 (*Pdcd1*)-expressing CD8^+^ T cells which are the target of PD-L1–PD-1 checkpoint blockade. These include a major *Prf1^high^Gzm^high^Pdcd1*^+^*Havcr2*^+^ cluster resembling “exhausted” T (TEX) cells and a *Pdcd1*^+^*Tcf7*^+^*Slamf6*^+^cluster resembling “stem-like” T cells^39,40^ (**Fig. 5A**, **Extended Data Fig. 8C-D**), that were predominantly tumour-resident and expanded with anti-PD-L1 (**Fig. 5B**, **Extended Data Fig. 8E**). *Pdcd1^+^* cells were evident among the cycling_CD8T cluster (**Extended Data Fig. 8F-G**) and increased in proliferation following anti-PD-L1, potentially underpinning their increased number (**Extended Data Fig. 8H-I**). Of note, the TEX cluster showed the largest transcriptional response to anti-PD-L1 (**Fig. 5C**), including upregulation of genes involved in “*TCR signalling*” and *“IL2-STAT5 signalling”* with potential anti-tumour benefits (**Extended Data Fig. 8J**). Indeed, Kaede-red^+^PD-1^+^ CD8^+^ T cells showed a significant increase in Granzyme B and IFNγ protein expression following anti-PD-L1 treatment (**Extended Data Fig. 8H,K-L**).

**Figure 5.**
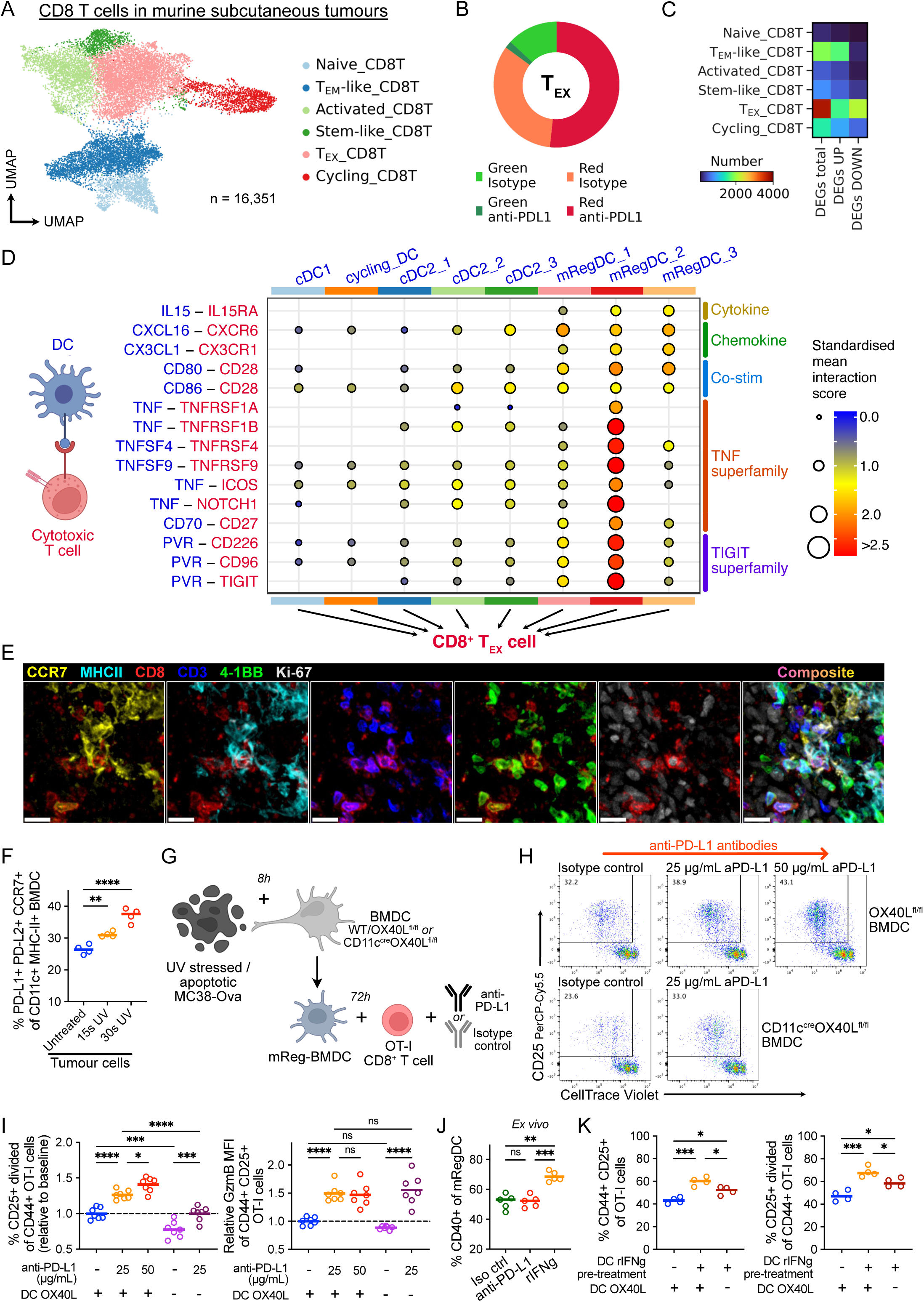
anti-PD-L1 promotes activating mRegDC-cytotoxic CD8^+^ T cell interactions in the TME. (A) UMAP of scRNA-seq of CD8^+^ T cells, from FACS-sorted CD45^+^ Kaede-green/red TILs 48h after photoconversion of subcutaneous MC38-Ova tumours. (B) Proportions of CD8^+^ TEX cells by Kaede profile and treatment. (C) Number of significant DEGs in anti-PD-L1 versus isotype control-treated tumours. DEGs calculated using Wilcoxon rank-sum test. (D) CellPhoneDB ligand-receptor analysis between tumour DCs and CD8^+^ TEX cells. Only significant interactions (*p* < 0.05) shown. (E) Representative confocal microscopy images of MC38 tumours, showing co-localisation of CCR7^+^MHC-II^+^ mRegDCs and CD3^+^CD8^+^4-1BB^+^Ki-67^+^ T cells. Scale bar, 20 μm. (F-K) *In vitro* and *Ex vivo* cultures. (F) Phenotype of BMDCs following culture with UV-irradiated MC38-Ova tumour cells. (G) Experiment set-up: DC-T cell co-culture. (H) Representative flow cytometry of OT-I activation and proliferation following co-culture with MC38-Ova-experienced mReg-BMDC; +/- anti-PD-L1, OX40L-expressing (+) or OX40L-deficient (-) BMDCs. (I) Quantification of (H) and GzmB expression. Values are plotted relative to cultures with isotype control antibodies and OX40L-expressing DCs (baseline), to facilitate comparisons across experiments. (J) Flow cytometry of mRegDCs from subcutaneously grown MC38-Ova tumours following 8h culture *ex vivo* with antibodies or recombinant IFNγ. (K) Flow cytometry of OT-I cells following co-culture with mReg-BMDC; DCs pre-treated with recombinant IFNγ (+) or PBS (-), OX40L-expressing (+) or OX40L-deficient (-) BMDCs. Data is representative of two independent experiments, one-way ANOVA with Šidák’s multiple comparisons test was used (F-K).

Given mRegDCs are a major source of PD-L1 in tumours, and the marked response of CD8^+^ TEX cells to anti-PD-L1 treatment, we sought to identify mRegDC-mediated interactions that might promote their activation and proliferation (**Fig. 5D**). Cell-cell communication analysis showed several previously described interaction pairs, including *Cxcl16* and *Il15* from CCR7^+^ DCs, which may recruit and sustain cytotoxic T cells (**Extended Data Fig. 9A**)^41^. Importantly, we identified TNF-superfamily interactions, including *TNFSF4*–*TNFRSF4* (OX40L–OX40), *TNFSF9*–*TNFRSF9* (4-1BBL–4-1BB), *CD70*–*CD27*, and *PVR*-mediated interactions upregulated between mRegDC_2, which were enriched with anti-PD-L1 treatment, and CD8^+^ TEX cells (**Fig 5D**). These TNF-superfamily ligands are known to promote T cell survival, proliferation and activation^35^, and PVR engages the activating receptor CD226, or its competing inhibitory receptor TIGIT, to control CD8^+^ T cell effector function^36^. We observed similar predicted interactions between mRegDC_2 and other PD-1-expressing CD8^+^ T cells, including *Tcf7^+^* stem-like cells (**Extended Data Fig. 9B**). Overall, mRegDC_3 had fewer predicted activating interactions with PD-1-expressing CD8^+^ T cells than mRegDC_1/2, supporting their status as an “exhausted” mRegDC state (**Extended Data Fig. 9C**), but all mRegDC states were a consistently high source of inhibitory PD-1 or CTLA-4 signals (**Extended Data Fig. 9D**).

To visualise the cellular interactions predicted by our analysis, we used IF microscopy to confirm that mRegDCs and CD8^+^ T cells spatially co-localise (**Extended Data Fig. 9E-F**). There were frequent interactions between tumour-residing mRegDCs and Kaede-red^+^CD3^+^CD8^+^ T cells, often within a perivascular niche, including proliferating (Ki-67^+^)4-1BB^+^ cytotoxic cells (**Fig. 5E**, **Extended Data Fig. 9G-I**), consistent with a role for mRegDCs in regulating anti-tumour cytolytic activity.

To test the functional importance of these predicted mRegDC-CD8^+^ T cell interactions, we generated mRegDC-like cells from bone marrow-derived dendritic cells (BMDC) by culturing with apoptotic MC38-Ova cells, as previously described^5^. This led to a robust expression of mRegDC markers, including PD-L1, PD-L2 and CCR7 (**Fig. 5F**, **Extended Data Fig. 10A-B**). Of note, the CCR7^+^PD- L2^+^ BMDCs (mReg-BMDC) generated by this system also expressed OX40L and PVR (**Extended Data Fig. 10C**), resembling the phenotype of tumour-retained mRegDC *in vivo*.

Co-culture of tumour-antigen (Ova)-experienced mReg-BMDC with naïve Ova-specific CD8^+^ T cells (OT-I) for 3 days (**Fig. 5G**) led to OT-I proliferation, which was not observed unless DCs were exposed to Ova (**Extended Data Fig. 10D-E**). Notably, the addition of anti-PD-L1 antibodies to the mReg- BMDC:OT-I co-culture resulted in an increase in activated (CD44^+^CD25^+^), clonally-expanded OT-I cells, and enhanced the production of granzyme B among activated cells (**Fig. 5H-I**, **Extended Data Fig. 10F**).

To assess the role of *Tnfsf4* (OX40L), a ligand expressed by the activated mRegDC_2 state increased by anti-PD-L1 treatment, and enables interactions with CD8^+^ T cells via OX40 (**Fig. 5D**), we generated OX40L-deficient mReg-BMDC from CD11c^cre^ *Tnfsf4*^fl/fl^ mice. OT-I cells co-cultured with OX40L- deficient mReg-BMDC showed reduced activation and proliferation (**Fig. 5H-I**, **Extended Data Fig. 10G-H**), suggesting that the OX40L:OX40 axis may be important for mRegDC function *in vivo*.

Finally, we asked whether anti-PD-L1 treatment directly influences the tumour mRegDC state, or whether this is driven by DC-extrinsic factors. For example, immune checkpoint therapy increases interferon gamma (IFNγ) production by PD-1^+^CD8^+^ T cells that co-localise with mRegDCs in tumours (**Extended Data Fig. 8K**), which may activate tumour DCs^32^. Administration of anti-PD-L1 antibodies to isolated, tumour antigen-experienced BMDC *in vitro* did not alter their phenotype, but the addition of recombinant IFNγ upregulated expression of OX40L, PVR and CD40, consistent with the mRegDC_2 state (**Extended Data Fig. 10I**). Tumours cultured *ex vivo* with IFNγ resulted in similar activation of mRegDCs (**Fig. 5J**). To assess the significance of IFNγ-induced changes, we pre-treated mReg-BMDC with IFNγ, prior to OT-I co-culture. This increased the activation and expansion of OT-I cells, but was reduced in cultures containing OX40L-deficient mReg-BMDC (**Fig. 5K**, **Extended Data Fig. 10J**). Altogether, these data support the conclusion that tumour mRegDC states can be manipulated to promote antigen-specific CD8^+^ T cell responses.

### Conserved mRegDC heterogeneity and CD8^+^ T cell crosstalk in human cancers

We asked if the time, tissue, and treatment-associated heterogeneity in mRegDCs observed in our murine model was pertinent to human cancers. In mRegDCs from human CRC (**Extended Data Fig. 1B**), there was a gradient of MHC-II expression (**Fig. 6A**). Importantly, markers upregulated on tumour-residing *MHC-II*^low^ mRegDCs in murine tumours were also preferentially expressed in the human *MHC-II*^low^ mRegDCs, including *IL15*, *PVR*, *TNFSF4*, *TNFSF9*, and *CD70*.

**Figure 6.**
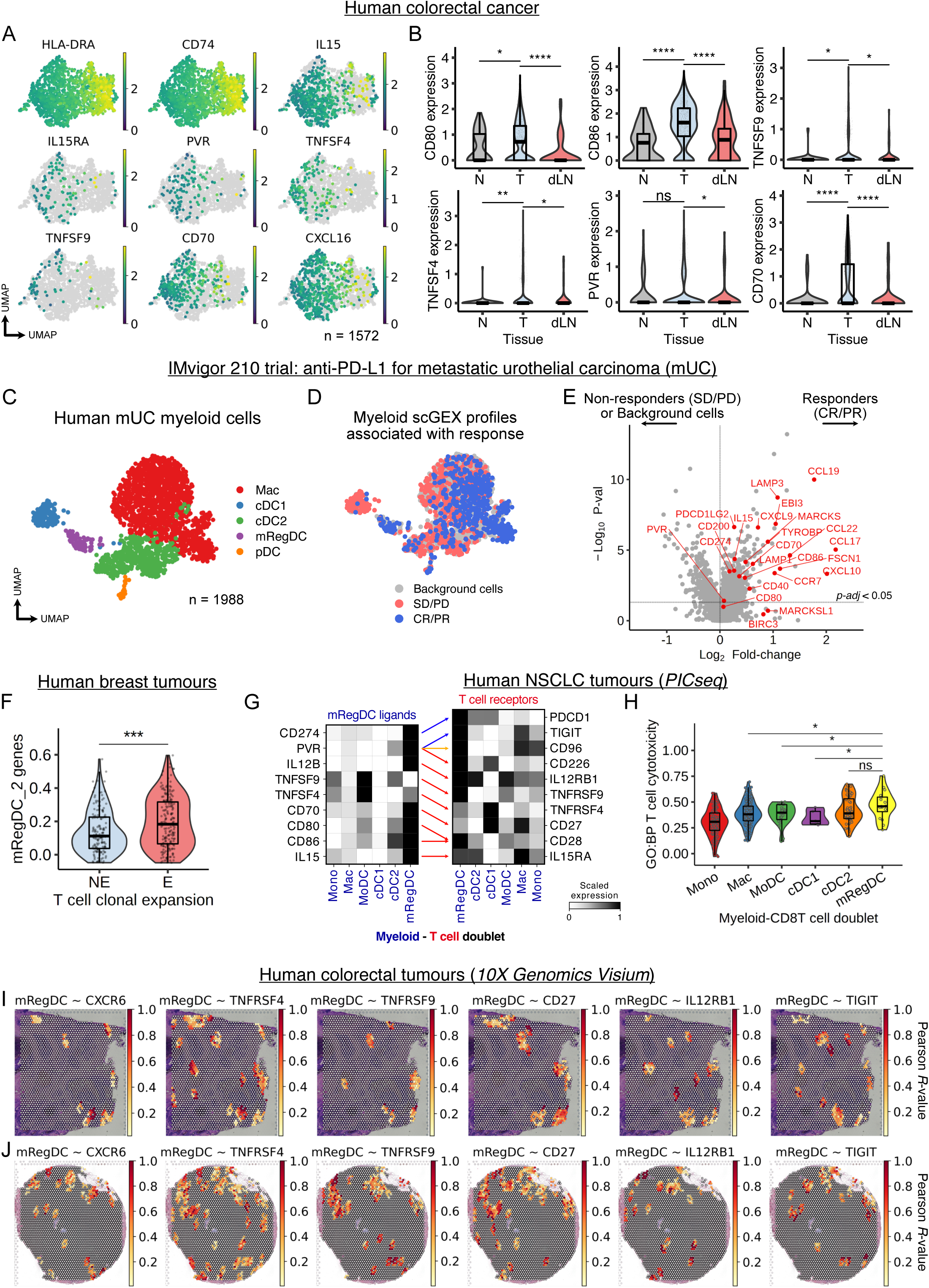
Conserved mRegDC heterogeneity and CD8^+^ T cell crosstalk in human cancer. (A) UMAP of scRNA-seq of mRegDCs from human CRC (GSE178341, *n* = 62 patients). (B) Expression of selected ligands in mRegDCs between tumour (T), normal adjacent tissue (N) and tumour-dLN from independent scRNA-seq data of human CRC (syn26844071, *n* = 63 patients). Wilcoxon rank-sum test was used. (C) UMAP of myeloid cells from scRNA-seq of human metastatic urothelial carcinoma (mUC, HRA000212, *n* = 11 patients). (D) Cells associated with responders (complete responder, CR; partial responder, PR) or non- responders (stable disease, SD; progressive disease, PD) in the IMvigor 210 trial (atezolizumab) for treatment of mUC; Scissor integration of tumour bulk RNA-seq samples from IMvigor210 (EGAS00001004343, *n* = 208 patients) and scRNA-seq from (C). (E) Differential gene expression in myeloid cells associated with responders (CR/PR) versus non-responders (SD/PD) from (D). (F) Gene signature scores for mRegDC_2 transcripts in mRegDCs from human breast cancer treated with anti-PD-1 antibodies. Comparison between T cell clonotype expanders (responders, *n* = 9 patients) versus non-expanders (non-responders, *n* = 20 patients, EGAS00001004809). Wilcoxon rank-sum test was used. (G) PICseq (myeloid-T cell doublets) of human NSCLC; expression of mRegDC-T cell ligand-receptor pairs, grouped by myeloid cell identity in each myeloid-T cell doublet (GSE160903, *n* = 10 patients). (H) Gene signature scores for “*GO:BP T cell mediated cytotoxicity (GO:0001913)*” (HLA genes removed), in myeloid-T cell PICs grouped by the myeloid identity. Wilcoxon rank-sum test was used. (I-J) Spatial correlation (Pearson R-value) of mRegDC signature scores and selected mRegDC-ligand receptors expressed by CD8^+^ T cells, in spatial transcriptomics (10X Genomics Visium) of independent human CRC tumour sections (*n* = 2).

To assess tissue-associated differences in mRegDCs, we analysed an independent human CRC dataset with paired scRNA-seq of tumour, dLN and normal adjacent tissue (**Extended Data Fig. 11A-C**)^42^. Consistent with our findings, there was no difference in *CCR7* expression, but *CD80*, *CD86*, *TNFSF4*, *TNFSF9*, *CD70* and *PVR* were higher in tumour-residing mRegDCs versus dLN and normal tissue, while *ICOSLG* was enriched in the dLN (**Fig. 6B**, **Extended Data Fig. 11D-E**). Therefore, the heterogeneity of mRegDCs in murine tumours are paralleled in human CRC.

Next, we sought to address whether intra-tumour mRegDCs, and their precise phenotype, influences clinical response to ICB. Atezolizumab (anti-PD-L1 antibody) is widely used in the treatment of metastatic urothelial carcinoma (mUC), but its efficacy is variable^43^. We analysed 208 bulk transcriptomes of tumour biopsies from the IMvigor210 trial for mUC^44,45^. In both responders and non-responders, mRegDC gene expression was positively correlated with effector CD8^+^ T cell transcripts, and was higher in responders (**Extended Data Fig. 11F**). To leverage the clinical response data from this cohort, we integrated the bulk transcriptomes with scRNA-seq of myeloid cells in mUC^46^ (**Fig. 6C**) using *Scissor*^47^, which enables identification of clinically relevant cell subpopulations and gene expression profiles. This analysis revealed that mRegDCs were enriched in responders (**Fig. 6D**). Indeed, myeloid cells associated with a favourable clinical response expressed transcripts associated with tumour-residing mRegDCs, including *CCR7*, *CD274*, *IL15*, *PVR* and *CD70* (**Fig. 6E**).

Moreover, in scRNA-seq of breast cancer (**Extended Data** Fig. 1D, **11G**)^24^, mRegDCs from tumours that successfully underwent T cell clonotype expansion following treatment with anti-PD-1 antibodies were enriched in transcripts associated with our ICB-induced mRegDC_2 cluster, versus non-responders (**Fig. 6F**). mRegDCs from T cell clonotype expanders also upregulated genes contributing to ‘*interferon gamma response*’, ‘*FcγR-mediated phagocytosis*’, ‘*lymphocyte co-stimulation*’, etc. (**Extended Data Fig. 11H**). Altogether, data from these treatment cohorts support the importance of an ICB-activated tumour-residing mRegDC state.

Next, we asked if the molecular crosstalk between mRegDCs and CD8^+^ T cells in mice is conserved in humans. In PIC-seq data from NSCLC^38^, we found that doublets containing mRegDCs highly expressed ligands we identified in murine tumours (**Fig. 5D**), and their corresponding T cell receptors were highly expressed in the same mRegDC-T cell conjugates (**Fig. 6G**). Moreover, mRegDC-CD8^+^ T cell doublets were most enriched for T cell cytotoxicity genes and ‘*TNFα response via NFκΒ*’ compared to other myeloid-CD8^+^ T cell combinations (**Fig. 6H**, **Extended Data Fig. 11I-J**).

Finally, we re-examined hotspots of mRegDC and effector CD8^+^ T cell co-localisation in spatial transcriptomics of CRC, breast cancer and melanoma (**Fig. 4D-F**). Molecules involved in mRegDC- CD8^+^ T cell crosstalk were expressed in these voxels, with spatial correlation of mRegDC-CD8^+^ T cell ligand-receptor pairs in all 3 tumour types (**Fig. 6I-J**, **Extended Data Fig. 12A-C**). Altogether, these data show that mRegDCs are critically positioned to regulate anti-tumour cytolytic activity, including via TNF-superfamily ligands such as OX40L, and PVR.

## Discussion

In this study, we combined scRNA-seq with a photoconvertible murine tumour model to unravel the spatio-temporal dynamics of tumour mRegDCs. Contrary to current assumptions, mRegDCs are heterogeneous, including sub-populations that either primarily contribute to LN migration or are retained in the tumour. Since CCR7^+^ DCs do not uniformly and instantaneously migrate to the dLN, and influence local tumour immunity, this activated DC subset cannot be unequivocally labelled as “migratory DC”. We found that tumour-retained mRegDCs acquire transcriptional features consistent with functional “exhaustion” with prolonged tumour residence, but following anti-PD-L1 treatment, are skewed towards a state enriched in T cell stimulatory molecules, capable of augmenting anti-tumour cytotoxic T cell responses. Their heterogeneity, crosstalk with CD8^+^ T cells, and association with response to ICB was conserved across human cancers.

Our work suggests that mRegDCs serve as a critical cellular immunoregulatory hub within the TME, through the provision of chemotactic, survival, activating, and inhibitory factors. We identified an mRegDC-CD8^+^ T cell axis, including specific interactions which control anti-tumour cytolytic activity. Consistent with this, acquisition of the full effector CD8^+^ T cell program requires engagement of CD80^high^ DCs in the local TME^48^, which we propose are mRegDCs. Recent studies also report that mRegDCs may engage other immune cells in tumours, including CXCL13^+^ CD4^+^ T cells, regulatory T cells and NK cells ^38,49–51^, and may reside in TLS^12,15^. How heterogeneous mRegDC states influence the survival or activation of other immune cells, including within TLS, warrants further investigation.

The success of ICB have led to combination immunotherapies^52^, exemplified by the recent success of tiragolumab (anti-TIGIT) + atezolizumab for NSCLC^53^. In CD8^+^ T cells, mechanistic convergence of PD-1 and TIGIT inhibitory pathways powerfully regulates cytotoxic function^54^. We found that mRegDCs are the major suppliers of both PD-1 and TIGIT ligands to CD8^+^ T cells, and importantly, there was an increase in PVR expression following anti-PD-L1 treatment, potentially regulating T cell activation via CD226/TIGIT engagement^36^. Independently, combining OX40 or 4-1BB agonists with anti-PD-L1 or anti-PD-1 antibodies respectively improved treatment outcomes in pre-clinical cancer models^55,56^. Given their diverse ligand profile, our data suggest that mRegDCs are a molecular hub through which these combinatorial immunotherapies synergise.

Several questions remain; While provision of PD-1 ligands by DCs are essential for effective ICB^16,17^, it remains unclear whether anti-PD-L1 antibodies directly alter DC function. Interestingly, ‘reverse signalling’ through PD-L1 has been described^57^, including effects on DC migration^58^. Our *in vitro* data does not support a DC-intrinsic effect of anti-PD-L1 on tumour DC phenotype, but a cell type-specific deletion of PD-L1 *in vivo* would be needed to definitively address this, as *in vitro* systems may not fully recapitulate the complexity of the TME. Next, knowledge of mRegDC ontogeny remain incomplete^6,8^, particularly the relative contribution of cDC subsets or monocytes to heterogeneous mRegDC states, which may influence their ability to support CD8^+^ versus CD4^+^ T cell responses. Our analyses suggest both cDC1 and cDC2 precursors contribute, but fate mapping experiments would be required to definitively prove this. Finally, whether the retention of mRegDCs in tumours is due to the acquisition of aberrant trafficking behaviour or due to intratumoral interactions, such as chemoattraction to stromal cells or tumour cells expressing CCR7 ligands^59,60^ remains to be resolved. Indeed, the presence of mRegDC in TLS suggest that local cues facilitate their retention.

Previous studies have drawn conflicting conclusions on the role of mRegDCs in tumours; Loss of TIM- 3 in DCs prevents the acquisition of the mRegDC programme, but facilitates maintenance of the effector CD8^+^ T cell pool^61^. Conversely, CXCL16 and IL-15 expressing CCR7^+^ DCs may recruit and sustain CXCR6^+^ cytotoxic T cells crucial in cancer immunosurveillance^41^. Our data help to resolve these contradictions; We propose that tumour mRegDCs are heterogeneous, with subset-specific capacity to both support and inhibit the cytotoxic T cell niche, presenting new opportunities for intervention.

## Methods

### Mice

Transgenic C57BL/6 Kaede, BALB/c Kaede, OX40L^+/Human-CD4^ reporter, OX40L^fl/fl^, and CD11c^cre^ OX40L^fl/fl^ mice are maintained and bred at the University of Birmingham Biomedical Services Unit. Wild-type C57BL/6 mice were maintained and bred at the University of Birmingham Biomedical Services Unit or the University of Cambridge Biomedical Services Gurdon Institute animal facilities. Mice were culled between the ages of 8 and 14 weeks. All animal experiments were conducted in accordance with Home Office guidelines and were approved by the University of Birmingham Animal Welfare and Ethical Review Body or the University of Cambridge Animal Welfare and Ethics Review Board. Mice were housed at 21°C, 55% humidity, with 12 h light-dark cycles in 7-7 individually ventilated caging with environmental enrichment of plastic houses plus paper bedding.

### Mouse subcutaneous tumour model

MC38 (kindly provided by Dr. Gregory Sonnenberg; Weill Cornell Medicine, New York, NY), CT26 (kindly provided by Professor Tim Elliot, University of Oxford, Oxford, UK) and MC38-Ova (obtained from AstraZeneca) murine colon adenocarcinoma cells were cultured in DMEM and supplemented with 2mM L-glutamine (Thermo Fisher Scientific), 10% FBS (F9665; Sigma-Aldrich), and penicillin-streptomycin (Sigma-Aldrich) at 37°C with 5% CO2. Cells grown in the log-phase were harvested and resuspended to 2.5 × 10^6^ cells/ml in Dulbecco’s PBS (Sigma-Aldrich) for tumour injection.

2.5 × 10^5^ tumour cells in 100 μl were subcutaneously injected into mice in the pre-shaved left flank area under anaesthesia via 2% gaseous isoflurane. For experiments involving anti-PD-L1 treatment, including scRNA-seq, MC38-Ova tumours were used; for DC phenotyping, specific tumour cells used for each experiment are included in figure legends. Tumour size was periodically measured with a digital Vernier calliper, and the volume was calculated using the formula V = 0.5 × a × b^2^ in cubic millimetres, where a and b are the long and short diameters of the tumour respectively. Tumour weights were measured at the endpoint of the experiment. Mice were sacrificed on day 13, 14, 15, or 16 (5h, 24h, 48h, 72h post-photoconversion respectively, where performed) and tumours were harvested for analysis. Where indicated for specific experiments, tumour-dLNs (i.e. left inguinal LN) and contralateral non-draining lymph nodes were also harvested.

### Administration of anti-PD-L1 antibodies

Anti-PD-L1 mouse IgG1 (Clone 80, SP16-260; AstraZeneca) or NIP228 isotype control mouse IgG1 (SP16-017; Astra Zeneca) were administered on day 7, 10, and 13 after tumour injection by intraperitoneal injection. Each dose consisted of 200 μg antibodies diluted in 200 μl PBS (10 mg/kg body weight). Tumour volume was measured on day 7, 10, 13 and the experiment endpoint.

### Labelling of tumour compartment by photoconversion

Photoconversion was performed as previously described^19,28^. Briefly, on day 13 after tumour injection, the subcutaneous tumour was exposed to a 405-nm wavelength focussed LED light (Dymax BlueWave QX4 outfitted with 8mm focussing length, DYM41572; Intertronics) for 3 minutes, with a 5-second break every 20 seconds, at a fixed distance of 1 cm. Black cardboard was used to shield the remainder of the mouse. We previously showed that this method resulted in complete conversion (99.9%) of host cells within the tumour from the default green fluorescence of the Kaede protein (Kaede-green) to the altered red fluorescence (Kaede-red), and that the cells in the dLN were fully protected from tumour photoconversion^19^. Of note, while converted cells express Kaede-red fluorescence, they also retain a weak Kaede-green signal. Overall, this enabled the discrimination of newly infiltrating (Kaede-green) and resident (Kaede-red) cell populations in the tumour, or cells that have egressed the tumour (Kaede- red) by their fluorescence profile. Moreover, using i.v. administration of anti-CD45 antibodies prior to culling, we previously showed that majority of cells (>95%) were in tumour tissue and were not intravascular contaminants^19^.

### Confocal microscopy and analysis

Tumour or LN samples were fixed in 1% paraformaldehyde (Electron Microscopy Services) for 24h at 4°C followed by 12h in 30% sucrose in PBS. 20μm sections were permeabilized and blocked in 0.1M TRIS, containing 0.1% Triton (Sigma), 1% normal mouse serum, 1% normal rat serum and 1% BSA (R&D Systems). Samples were stained for 2h at RT in a humid chamber with the appropriate antibodies, listed in Supplementary table 1, washed 3 times in PBS and mounted in Fluoromount-G® (Southern Biotech). Images were acquired using a TCS SP8 (Leica microsystems, Milton Keynes, UK) confocal microscope. Raw imaging data were processed using Imaris v9.7.2 (Bitplane).

Iterative staining of sections was performed as previously described^62^. Samples were prepared and stained as detailed above. Following acquisition, the coverslips were removed, and slides were washed 3 times in PBS to remove residual mounting medium. Bleaching of the fluorochromes were achieved by submerging the slide in a 1mg/mL solution of lithium borohydride in water (Acros Organics) for 15 minutes at room temperature. The slides were then washed 3 times in PBS prior to staining with a different set of antibodies. The process was repeated twice. Raw imaging data were processed using Imaris using CD31 as fiducial for the alignment of acquired images.

For quantification and co-localisation analysis, processed fluorescence imaging data from Imaris were further analysed using QuPath^63^. Hoechst nuclear staining was first used to perform automated cell detection with the nucleus diameter setting at 3-10 μm. Thereafter, detections were manually annotated to identify 100 cells each for mRegDCs or CD8^+^ T cells per image. mRegDCs were annotated based on CD45^+^ MHC-II^+^ CCR7^+^ staining and dendritic morphology. CD8^+^ T cells were annotated based on CD45^+^ CD3^+^ CD8^+^ CCR7^+/-^ staining and spherical morphology. Vessels (CD31^+^) and tumour or stromal cells (CD45^-^) were annotated as detections to ‘ignore’. The manual annotations were used to train a semi-automated random trees object classifier, to automate annotation of remaining cells, using the following parameters: nuclear and cellular morphology, CD45, CD3, CD8, MHC-II, CCR7, and CD31 staining intensity. The classification output and centroid positions for all cell detections were exported for further analysis in R. For correlation analysis, tumour sections were divided into grids approximately 20-30 cell detections wide (200 μm). Number of cell detections of each class were counted per grid and Pearson correlation was applied to quantify spatial co-localisation.

### Tissue dissociation

Tumours were cut into small pieces using surgical scissors, and incubated with 1 mg/ml collagenase D (Roche) and 0.1 mg/ml DNase I (Roche) in a volume of 1.2 ml RPMI media at 37°C on a thermomixer (Eppendorf) for 20 min; or tumours were digested using Tumour Dissociation Kit (Miltenyi Biotec) and gentleMACS Dissociator (Miltenyi Biotec) for 40 minutes at 37°C according to the manufacturer’s protocol. The gentleMACS protocol was used for scRNA-seq experiments. Subsequently, the sample was filtered through a 70 μm strainer to remove undigested tissue debris. Next, dead cells were removed using Dead Cell Removal Kit and LS Columns (Miltenyi Biotec), according to the manufacturer’s instructions. Lymph nodes were cleaned and dissected in RPMI 1640 medium (Thermo Fisher Scientific) and crushed through a 70 μm strainer. Thereafter, cells were centrifuged at 400 *g* at 4°C for 5 min and resuspended in FACS staining buffer (2% FBS; 2mM EDTA in PBS) for flow cytometry.

### Flow cytometry

Cell suspensions were subjected to Fc block with anti-CD16/32 (BioLegend) diluted in FACS staining buffer on ice for 15 min before staining with surface markers, listed in Supplementary table 1, diluted in FACS staining buffer on ice for 30 min, and subsequently, a live/dead stain. Where applicable, cells were then fixed with eBioscience intracellular fixation buffer (Thermo Fisher) for 30 min and stained for intracellular markers diluted in eBioscience permeabilization buffer (Thermo Fisher) at 4°C overnight. 1 x 10^4^ counting beads (Spherotech) were added to stained samples at the final step, to calculate absolute cell numbers. Data were acquired on the LSR Fortessa X-20 (BD) using FACSDiva v8.0.2 software (BD) or CytoFLEX (Beckman Coulter) using CytExpert v2.5 (Beckman Coulter) and analysed with FlowJo v10.8.1 (BD).

### Single-cell isolation

MC38-Ova tumours were injected in age-matched female Kaede C57BL/6 mice, treated with anti-PD- L1 antibodies, and photoconverted on day 13 as described above. Mice with tumours of similar sizes were collected 48h after tumour photoconversion. After tumour digestion, as described above, cell suspensions were stained for CD45 BV786, TER119 PE-Cy7, CD11b BV421, NK1.1 BV650, Live/dead APC-Cy7 on ice for 30 min (Supplementary table 1). Subsequently, cells were centrifuged at 400 *g* at 4°C for 5 min and resuspended in FACS staining buffer for sorting. Tumour-infiltrating lymphocytes (TIL; Live CD45^+^TER119^-^Kaede^+^CD11b^-/low^NK1.1^low/hi^) and tumour-infiltrating myeloid cells (Live CD45^+^TER119^-^Kaede^+^CD11b^+^NK1.1^-^) from anti-PD-L1 or isotype-control treated tumours were sorted with a FACS Aria II Cell Sorter (BD) into two groups per cell type, based on the presence or absence of Kaede-red signal. CD45^+^ cells were only sorted to ‘myeloid’ or ‘TIL’ (CD11b^+^ or CD11b^-/low^ respectively) fractions to ensure appropriate representation of various cell types in the scRNA-seq data. All single cell transcriptomes were combined at the analysis stage, before cell type annotation, to ensure all CD45^+^ immune cells, regardless of surface CD11b or NK1.1 expression, are represented in the final analysis and annotated based on their transcriptome.

Tumour-dLNs from mice treated with the same experimental protocol were harvested and digested as described in the preceding paragraph. Cell suspensions were stained for CD45 BV785, TER119 PE- Cy7, Live/dead APC-Cy7 on ice for 30 min. Live CD45^+^Ter119^-^Kaede^+^Kaede-red^+^ cells were sorted, to identify immune cells that have migrated from photoconverted tumours (Kaede-red) to the tumour- dLNs. Inguinal lymph nodes from control mice (no tumour cells injected) were also processed for scRNA-seq to obtain a representation of homeostatic cell populations in the LN. This was done to facilitate accurate annotation and integration of Kaede-red cells in the tumour-dLN, where specific populations arriving from the tumour would be disproportionately over-represented, with the LN cellular landscape. Control LN cell suspensions were stained for B220 BV421, CD11c BV786, TER119 PE-Cy7, Live/dead APC-Cy7 on ice for 30 min. Thereafter, a 3-way sort was performed consisting of Live Kaede^+^TER119^-^CD11c^+^B220^-^ (myeloid), Kaede^+^TER119^-^CD11c^-^B220^+^ (B cells), and Kaede^+^TER119^-^CD11c^-^B220^-^ fractions. The three subsets were then mixed at a ratio of 1:1:1 to generate a cellular suspension enriched for DCs and with reduced frequency of B cells. scRNA-seq of control LNs and tumour-dLNs were sequenced and analysed together.

### Single-cell library construction and sequencing

Gene expression libraries from were prepared from FACS-sorted populations of single cells using the Chromium Controller and Chromium Single Cell 3’ GEM Reagent Kits v3 (10x genomics, Inc.) according to the manufacturer’s protocol. The resulting sequencing libraries comprised of standard Illumina paired-end constructs flanked with P5 and P7 sequences. The 16 bp 10x barcode and 10 bp UMI were encoded in read 1, while read 2 was used to sequence the cDNA fragment. Sample index sequences were incorporated as the i7 index read. Paired-end sequencing (2 x 150 bp) was performed on the Illumina NovaSeq 6000 platform. The resulting .bcl sequence data were processed for QC purposes using bcl2fastq software (v2.20.0.422) and the resulting fastq files were assessed using FastQC (v0.11.3), FastqScreen (v0.9.2) and FastqStrand (v0.0.5) prior to alignment and processing with the CellRanger (v6.1.2) pipeline.

### Processing of scRNA-seq

Single-cell gene expression data from CellRanger count output (filtered features, barcodes, and matrices) were analysed using the Scanpy^64^ (v1.8.2) workflow. Raw count data from the myeloid and TIL sorts were concatenated. Doublet detection was performed using Scrublet^65^ (v0.2.1), with cells from iterative sub-clustering flagged with outlier Scrublet scores labelled as potential doublets. Cells with counts mapped to >6000 or <1000 genes were filtered. The percentage mitochondrial content cut- off was set at <7.5%. Genes detected in fewer than 3 cells were filtered. Total gene counts for each cell were normalised to a target sum of 10^4^ and log1p transformed. This resulted in a working dataset of 80,556 cells. Next, highly variable features were selected based on a minimum and maximum mean expression of ≥0.0125 and ≤3 respectively, with a minimum dispersion of 0.5. Total feature counts, mitochondrial percentage, and cell cycle scores, where indicated, were regressed. The number of principal components used for neighbourhood graph construction was set to 50 initially, and subsequently 30 for myeloid and TIL subgroup processing. Clustering was performed using the Leiden algorithm with resolution set at 1.5 for initial annotations, but subsequently sub-clustering was performed at lower resolutions (0.7-1.0) for analysis of myeloid cell, DC, T cell, and CD8^+^ T cell subsets. Uniform manifold approximation and projection (UMAP, v0.5.1) was used for dimensional reduction and visualisation, with a minimum distance of 0.3, and all other parameters according to the default settings in Scanpy.

### Analysis of scRNA-seq from mouse tumour models

Broad cell types of interest were subset (eg. DCs from myeloid cells) and re-clustered as described above. Resulting clusters were annotated using canonical marker gene expression and published transcriptomic signatures. Unless otherwise indicated, log-transformed expression values were used for plotting. Gene set scoring was performed using Scanpy’s tl.score_genes tool. Gene sets were obtained from the Molecular Signature Database (MSigDB) inventory, specifically Hallmark, KEGG, or Gene Ontology (GO), using the R package msigdbr (v7.5.1) or published RNAseq data (Supplementary table 2). The original published mRegDC signature gene list^5^ was used for the initial scoring and identification of mRegDCs. Differential gene testing was performed using the Wilcoxon rank sum test implemented in Scanpy’s tl.rank_genes_groups. Analysis of MC38-Ova and B16 tumours and their dLNs followed these same methods. Gene regulatory network and transcription factor regulon activity analyses was performed in pyScenic (v0.12)^66^.

Trajectory analysis was performed using partition-based graph abstraction (PAGA)^67^ and the Palantir algorithm^68^. For Palantir pseudo-time analysis, the differentiation trajectory was rooted in the Kaede- green dominant cluster, which represents the cellular subset most associated with newly-infiltrating cells. RNA velocity analysis was performed using scVelo^69^, following the default pipeline. Integration and label transfer of lymph node DC scRNA-seq data and tumour DCs, or cycling CD8^+^ T cells and non-cycling CD8^+^ T cells was performed using Scanpy’s tl.ingest tool. For analysis of cycling CD8^+^ T cells, cell cycle regression was first performed on the isolated cluster using Scanpy’s pp.regress_out function, before re-integration.

Gaussian kernel density estimation to compute density of cells in the UMAP embedding was performed using Scanpy’s tl.embedding_density. Differential abundance analysis of *k*-nearest neighbour (kNN) defined cellular neighbourhoods was performed using Milo^70^. Spcifically, the kNN graph was constructed with a *k* parameter of 30 and initial random sampling rate of 0.1. Cellular neighbourhoods with constituent cells comprising less than 70% of a previously defined cluster were designated as mixed neighbourhoods. CellPhoneDB^71^ (v2.1.7) was used for cell-cell communication analysis, using the default parameters and normalised expression values as the input. CellPhoneDB output was visualised using ktplots (https://github.com/zktuong/ktplots).

For pathway analysis between Kaede-red versus Kaede-green or anti-PD-L1 versus isotype control groups in the scRNA-seq dataset, a pseudo-bulk approach was first applied to single-cell gene expression data (https://github.com/colin-leeyc/CLpseudobulk), to increase robustness for pathway analysis and overcome limitations associated with differential expression testing on single cells^72^. Briefly, filtered, raw count data (prior to normalisation, transformation, or scaling) from each condition were randomly sorted into artificial replicates, with iterative bootstrapping applied to random sampling (*n* >10). Pseudo-bulked count matrices were normalised using the median-of-ratios method, implemented in DESeq2^73^ (v1.34), and differential gene expression testing was performed using the Wald test. Pre-ranked gene set enrichment analysis^74^ (GSEA) was implemented in fgsea (v1.24), using the averaged Wald statistic over *n* >10 iterations of random sorting into pseudo-bulked replicates, as the gene rank metric. Leading edge genes were identified for further analysis. GO term over- representation analysis was implemented in topGO^75^ (v2.50), using the top 100 genes with the highest loadings for each principal component (PC) of interest, following PC analysis.

### Analysis of scRNA-seq from human tumours

scRNA-seq data from human solid tumours were downloaded from public repositories, which are listed under *Data availability* with references. Where necessary, data access permissions were sought and approved prior to download. Data was analysed using the Scanpy (v1.8.2) workflow, as outlined in the sections above. Filtering for quality control was performed according to the parameters outlined in the original publications. scRNA-seq integration was performed using the batch-balanced KNN (with ridge regression) approach or Harmony algorithm with sequencing batch as the batch term, where available and sufficient, or patient ID if sequencing batch was not available. Cell annotations from original publications were checked and refined, particularly where myeloid cell annotations were not complete or the focus of the original publication.

Myeloid cells were subset and used for further analysis. In total, this included re-analysis of 41,624 and 43,193 myeloid cells from two independent CRC datasets, 16,688 myeloid cells from breast tumours, 8,555 myeloid cells from cutaneous melanoma, 11,663 myeloid or T cells from NSCLC including 901 myeloid-T cell doublets, and 1,988 myeloid cells from metastatic urothelial carcinoma. mRegDCs were identified using expression of migratory transcripts and mRegDC signature genes^5^.

The only exception to the Scanpy workflow was for analysis of NSCLC PICseq data, where the Metacell workflow was utilised, as documented in the original publication, for analysis and annotation of single-cells^38^. This was to ensure compatibility with the PICseq deconvolution algorithm^76^, which was used to analyse sorted doublets. Gene expression raw count data from PICs only were obtained and input to the Scanpy workflow for visualisation and further analysis. Doublet deconvolution results for contributing myeloid cell or T cell identities in each PIC were retained.

Subsequently, analysis of gene expression profiles, gene signature scoring, cell-cell communication, were as described in earlier sections. For analysis of enrichment of mRegDC_2 genes in the breast cancer dataset, where anti-PD-1 response data was available, mRegDC_2 genes were first identified as DEGs between mRegDC_2 versus remaining mRegDCs (Wilcoxon rank-sum test) in scRNA-seq from murine tumours. 85 differentially upregulated genes (*p*-adj < 0.05, log2Fold-change > 1) were converted to equivalent human gene symbols using Ensembl and used to score mRegDCs from human breast cancer (Fig. 6F).

### Analysis of bulk transcriptomics

Bulk RNA-seq data were downloaded from public repositories, listed under *Data availability* with references. Where necessary, data access permissions were sought and approved prior to download. TCGA data was accessed via TCGAbiolinks, using STAR-aligned reads^23^. For bulk transcriptomics data from both TCGA and the IMvigor 210 trial, transcripts-per-million (tpm) normalised values were used for analysis. Cellular deconvolution was performed using xCell^77^, implemented in the webtool using default parameters. Gene signatures scoring was performed using single-sample GSEA^78^ (ssGSEA, v10.1.0), implemented in GenePattern, followed by normalisation (scaled between 0 and 1) of gene set enrichment scores. Cell type signatures for ssGSEA were derived from cell-specific differentially expressed genes in scRNA-seq of human cancers. Pearson correlation was used to assess for correlation of cellular proportions or cell-specific signatures of interests in biopsy samples. Survival analysis was performed on overall survival (months), using the median mRegDC signature score for stratification to ‘high’ and ‘low’ groups, and log-rank test was applied to survival curves. For integration of bulk transcriptomic data from the IMvigor210 trial with accompanying clinical response data with scRNA-seq of myeloid cells in mUC, the Scissor pipeline was used^47^. Scissor enables identification of single-cells and gene expression profiles within cell subpopulations that are significantly associated with phenotypes obtained from bulk expression data. A logistic regression model using the binary outcome of clinical responders (CR/PR) vs non-responders (SD/PD) was used, and the alpha parameter was set at 0.05, per default settings.

### Analysis of spatial transcriptomics

10x Visium spatial transcriptomics were analysed using the standard Scanpy (v1.8.2) workflow, using the default SpaceRanger outputs including spot alignments to corresponding tissue haematoxylin and eosin or immune-fluorescence images. Spots with fewer <500 counts, >80000 counts, and percentage mitochondrial content >15% were filtered. Genes detected in fewer than 5 spots were filtered. Total feature counts, mitochondrial percentage, and cell cycle scores were regressed out. Expression values plotted are log1p-transformed 10^4^-sum-normalised values. Gene set scoring was performed using Scanpy’s tl.score_genes tool. Calculation of spatial correlation was implemented using the correlationSpot function in ktplots (https://github.com/zktuong/ktplots), as previously described^79^. Briefly, k = 6 nearest neighbourhoods were extracted from a kNN graph computed from the spatial location of each Visium spot. Pearson correlation was performed on each neighbourhood using gene expression values or gene signature scores and averaged across the neighbourhoods. Correlation values were only returned if expression value or signature scores were detected in all spots, and above a significance threshold of *p*<0.05.

### Bone marrow derived dendritic cells

Bone marrow from wild-type C57BL/6, OX40L^fl/fl^, CD11c^cre^ OX40L^fl/fl^, or OX40L^+/Hu-CD4^ reporter mice were isolated by flushing femurs and tibias with RPMI, supplemented with 10% fetal calf serum, 1% penicillin/streptomycin, 1% L-glutamine and 1% sodium pyruvate (cRPMI). Bone marrow cells were strained through a 70 μm filter, centrifuged, and resuspended in RBC lysis buffer for 2 minutes on ice. Cells were plated in cRPMI in tissue-culture-treated 10 cm dishes. 20 ng/mL murine recombinant GM-CSF (Peprotech) and 5 ng/mL murine recombinant IL-4 (Peprotech) was added to generate bone-marrow derived dendritic cells (BMDC). Half of the medium was removed on day 2 of differentiation and new pre-warmed medium supplemented with GM-CSF and IL-4 (2X concentrations) were added. The culture medium was entirely replaced on day 3 with fresh warmed cRPMI + GM-CSF (20 ng/mL) only. On day 6, non-adherent cells in the culture supernatant were harvested for tumour cell-line co-culture.

### In vitro cultures

Ovalbumin-expressing MC38 (MC38-Ova) cells were plated tissue-culture-treated dishes in FCS-supplemented DMEM media (+ 5mM HEPES), as above. 24h after (80% confluence), the MC38-Ova monolayer was exposed to UV light (302 nm, 30s, 200 J m^-2^ s^-1^) to induce apoptosis, and further incubated for 24h. Day 6 BMDC were added to apoptotic MC38-Ova cells, and non-adherent cells were collected after 8h of co-culture for analysis by flow cytometry or further application. Either 1.5×10^5^ MC38-Ova cells were added to 24-well plates and 0.5×10^5^ BMDC were added 48h later, or 4.5×10^6^ MC38-Ova were added to 10 cm dishes and 1.5×10^6^ BMDC were added 48h later. Where indicated, 10 ng/mL murine recombinant IFNγ (Peprotech) or 25 μg/mL of anti-PD-L1 antibodies or isotype control antibodies were added to the tumour-DC cultures for 8h.

For OT-I T cell co-culture, 30s UV exposure was used to induce low-to-moderate MC38-Ova apoptosis for culture with BMDCs. Naïve CD8^+^ OT-I T cells (Live CD3^+^ CD8b^+^ CD62L^+^ CD44^-^) were FACS sorted from the spleens of OT-I × *Rag1*^-/-^ mice. Tumour antigen-experienced BMDC (Live CD45^+^ CD11b^+^ CD11c^+^ MHC-II^+^ PD-L2^+^) were FACS sorted from MC38-Ova tumour-DC cultures after 8h, as above. Sorted naïve OT-I T cells were labelled with CellTrace Violet (CTV, Thermo Fisher) at 37°C for 20 minutes, washed, and counted before use. 30,000 BMDC were incubated with 120,000 OT-I T cells in cRPMI in a 48 well tissue culture plate. For a positive control for OT-I stimulation, day 6 BMDCs were incubated with endotoxin-free ovalbumin protein (100 μg/mL) and LPS (20 ng/mL), before FACS sorting and culture with OT-I cells. For negative controls, BMDCs stimulated with apoptotic MC38 cells (no ovalbumin antigen), OT-I cultured with ovalbumin and LPS (no DC), or OT-I only were used. Anti-PD-L1 antibodies or isotype control antibodies were added on day 0 and day 2 as indicated. T cell activation and proliferation was assessed by flow cytometry after 3 days. For IFNγ pre-treatment of BMDC, recombinant IFNγ was added to tumour-DC cultures for 8h, washed twice to remove free cytokine, and FACS sorted, before culturing with OT-I cells. For analyses where data from multiple independent experiments were combined, values were normalised to respective replicates of cultures containing isotype control antibodies and OX40L-expressing DCs (baseline), to facilitate comparisons across experiments.

### *Ex vivo* cultures

For *ex vivo* tumour cultures, tumours were digested as described above, and centrifuged in 30% Percoll for 20 minutes to remove tissue debris, prior to dead cell removal. Tumour cell suspensions were resuspended in pre-warmed cRPMI and split equally into 3 wells for paired stimulation, to enable intra-tumour comparisons. *Ex vivo* cultures were performed at 37°C in 48w plates with 300 μL per well containing 10^7^ cells / mL. Isotype control or anti-PD-L1 antibodies (25 μg/mL each), or recombinant murine interferon gamma (10 ng/mL, Peprotech) was added for 8h. Cells were collected, culture plates washed twice with ice-cold PBS supplemented with 2mM EDTA and 10% FBS, and filtered before analysis by flow cytometry.

### Statistical analysis

Mice were gender-matched (all female). Tumour growth curves for anti-PD-L1 and isotype control treated groups were analysed using two-way ANOVA and Sidak’s multiple comparisons test and are presented as mean ± SEM. Analysis for RNA sequencing is as described above, including statistical frameworks used. Statistical tests were implemented for two principal purposes; to compare expression values (RNA or protein) between samples, or to compare proportions of defined cell subsets. All experiments were performed with biological replicates, and the specific statistical tests applied are indicated in the figure legends. All statistical tests applied were two-tailed. Paired statistical tests were used where different populations from the same animal were compared. All animal experiments were randomised prior to experimental intervention. Statistical significance are denoted as follows: ns, not significant; \**P* < 0.05; \*\**P* < 0.01; \*\*\**P* < 0.001; \*\*\*\**P* < 0.0001. Statistical analyses were performed in R or GraphPad Prism.

## Data and code availability

The scRNA-seq data have been deposited on the GEO public repository under accension numbers GSE221513 and GSE221064. Additional data or information will be made available upon reasonable request. Published data was accessed and downloaded from public GEO, SRA, EGA and Synapse repositories using the following ascension numbers: scRNA-seq of CRC (GSE178341 and syn26844071)^26,42^; scRNA-seq of breast cancer (EGAS00001004809)^24^; scRNA-seq of melanoma (GSE123139)^25^; scRNA-seq of mUC (HRA000212)^46^; scRNA-seq and PICseq of NSCLC (GSE160903)^38^; TCGA (https://portal.gdc.cancer.gov, via TCGAbiolinks)^23^; IMvigor210 bulk RNA-seq (EGAS00001004343)^44^; 10x Visium (https://www.10xgenomics.com/resources/datasets)^37^. All analysis performed are described under *Methods*. Source code for key softwares or computational pipelines used are referenced. Additional code used for analysis, as described above, have been made available on GitHub repositories.

## Author Contributions

CYCL designed and performed experiments, analysed data, performed computational analyses, and wrote the manuscript. BCK and ID performed experiments and analysed data. ZKT and TH provided project guidance. NR, FG, ZL, CW, SW and DP performed experiments. GC, SAH and SJD gave input and reviewed the manuscript. RR supported the project. DW conceived the project, designed experiments, provided guidance, and edited the manuscript. MRC conceived the project, designed experiments, provided guidance, and wrote the manuscript.

## Acknowledgments

We thank A. Ptasinska and other members of Genomics Birmingham at University of Birmingham for all their help with single-cell RNA-sequencing experiments, and the University of Birmingham Flow Cytometry Facility. We thank Dr. Y. Miwa (Tsukuba University, Tsukuba, Japan) and Dr. O. Kanagawa (RIKEN Research Center for Allergy and Immunology, Yokohama, Japan) and Dr. M. Tomura (Osaka Ohtani University, Tondabayashi, Japan) for the Kaede mice.

---

**Extended Data Figure 1.**
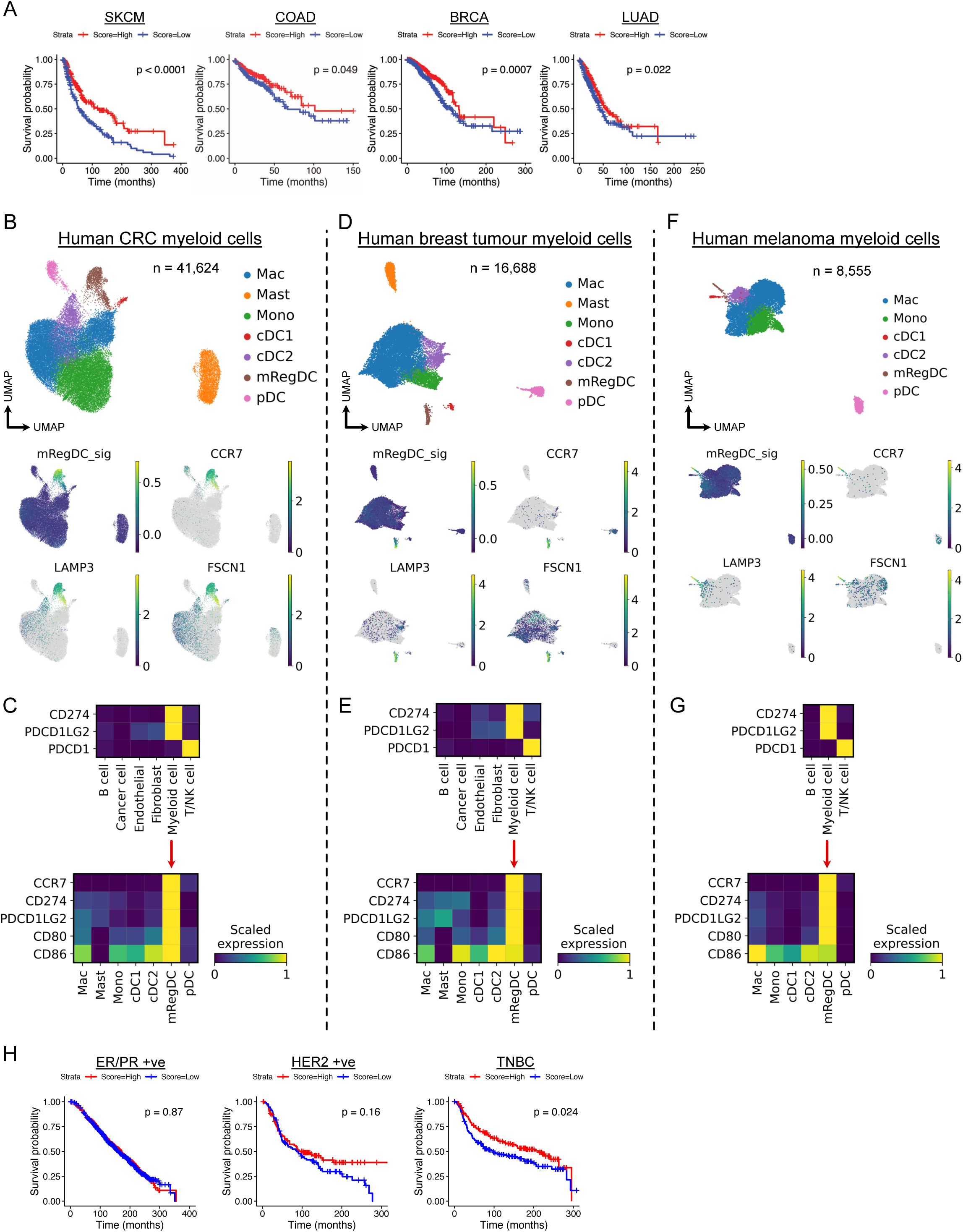
mRegDCs in human cancer. (A) Kaplan-Meier analysis of overall survival rate in skin cutaneous melanoma (SKCM, *n =* 473), colorectal adenocarcinoma (COAD, *n =* 521), breast invasive carcinoma (BRCA, *n =* 1226), and lung adenocarcinoma (LUAD, *n =* 598) from TCGA, stratified by enrichment of mRegDC signature genes. (B) Uniform manifold approximation projection (UMAP) of myeloid cells from scRNA-seq of human CRC (GSE178341, *n* = 62 patients) and expression of mRegDC genes (mRegDC signature score^5^, *CCR7*, *LAMP3*, *FSCN1*). (C) Expression of *CD274* (PD-L1), *PDCD1LG2* (PD-L2) and *PDCD1* (PD-1), *CCR7*, *CD80* and CD86 by cell-type in scRNA-seq of human CRC. (D) UMAP of myeloid cells from scRNA-seq of human breast cancer (EGAS00001004809, *n* = 29 patients) and expression of mRegDC genes. (E) Expression of selected genes by cell-type in scRNA-seq of human breast cancer. (F) UMAP of myeloid cells from scRNA-seq of human melanoma (GSE123139, *n* = 25 patients) and expression of mRegDC genes. (G) Expression of selected genes by cell-type in scRNA-seq of human melanoma. (H) Kaplan-Meier analysis of overall survival rate in hormone receptor-positive (oestrogen receptor, ER, or progesterone receptor, PR, *n* = 1369), human epidermal growth factor receptor 2 (HER2)-positive (*n* = 185), or triple-negative breast cancer (TNBC, *n* = 299) from METABRIC. *L*og-rank test was used.

**Extended Data Figure 2.**
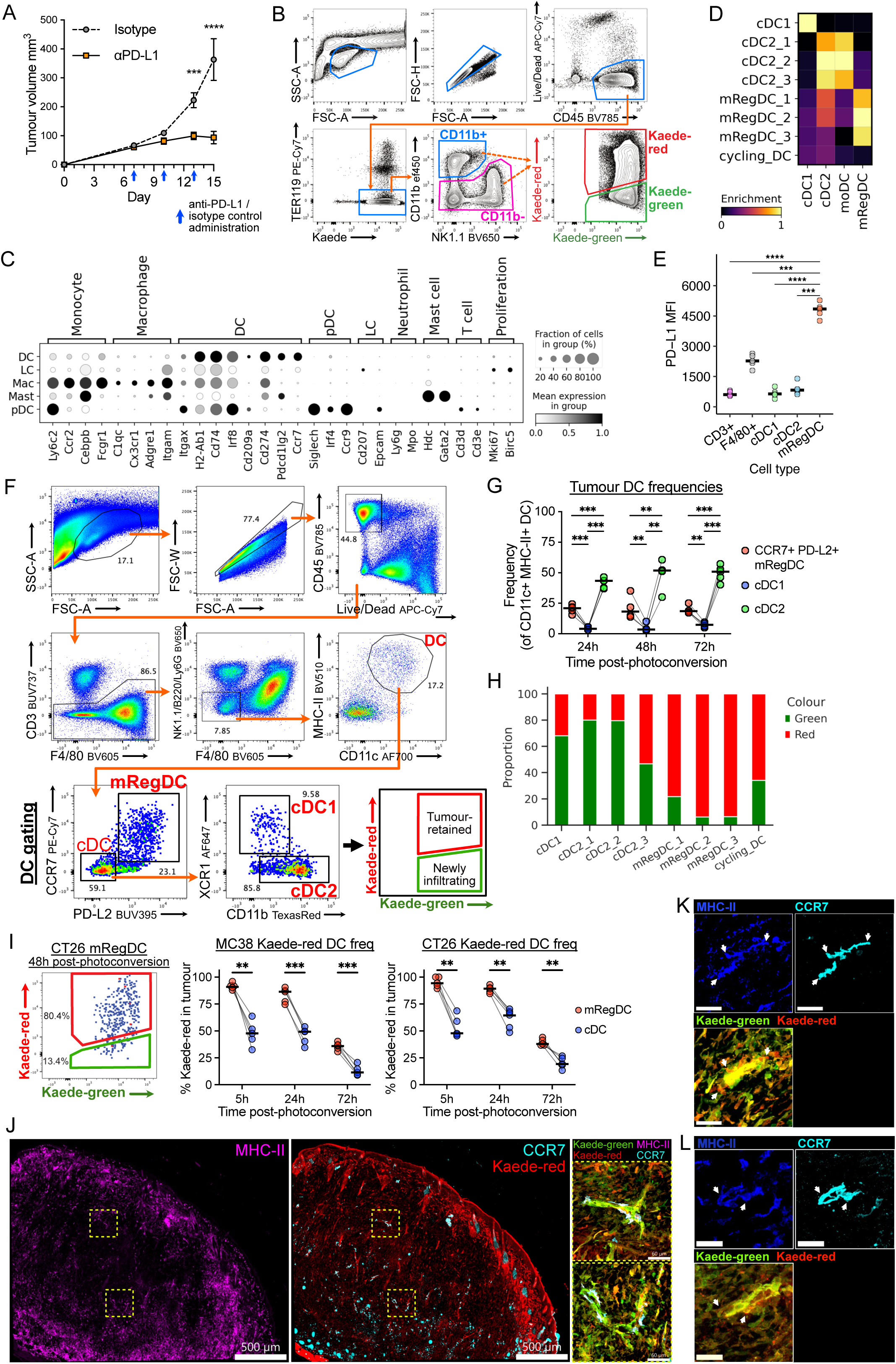
Single cell profiling of mouse subcutaneous tumours. (A) Tumour growth curves comparing anti-PD-L1 antibody-treated versus isotype control antibody treatment, administered on day 7, 10 and 13 in MC38-Ova tumours. Data shown is related to the scRNA-seq experiment (Fig. 1) and includes 5 mice per condition. (B) Fluorescence-activated cell sorting (FACS) strategy for isolation of tumour immune cells for scRNA-seq. CD45^+^ cells were only sorted to CD11b^+^ or CD11b^-/low^ fractions to ensure appropriate representation of various cell types in the scRNA-seq data, but both fractions were combined for scRNA-seq analysis and annotated based on their transcriptome. (C) Canonical marker gene expression in myeloid cell subsets, related to Fig. 1B. (D) Gene signature scores of publis hed transcriptomic signatures in DC clusters from Fig. 1C. Monocyte-derived DCs (moDC) signatures scored highly in cDC2 clusters, consistent with known challenges in distinction of cDC2 and moDC^7^. (E) Flow cytometry of surface PD-L1 expression in MC38-Ova tumours. (F) Representative flow cytometry gating strategy for tumour and LN DCs. mRegDCs were identified as Live, CD45^+^ lineage^-^ (CD3, NK1.1, B220, Ly6G, F4/80) CD11c^+^MHC-II^+^ PD- L2^+^CCR7^+^ cells; (PD-L2 expression was more restricted to mRegDCs than PD-L1.) Where Kaede transgenic mice were used, Kaede^+^ cells were gated. (G) Flow cytometry of DC composition in MC38-Ova tumours. Points represent independent mice. (H) Kaede fluorescence by DC cluster in the scRNA-seq data. (I) Flow cytometry of Kaede fluorescence in DCs from MC38 and CT26 tumours; photoconversion time course. Points represent independent mice. Paired t-test with FDR correction was used (G,H). (J) Representative microscopy of MC38 tumours 72h after tumour photoconversion. Insets highlight selected regions with Kaede-red CCR7^+^MHC-II^+^ DCs. (K-L) Zoomed-in representative microscopy of MC38 (K) and CT26 (L) tumours 72h after tumour photoconversion. Scale bar, 30μm; arrows, tumour-residing mRegDCs.

**Extended Data Figure 3.**
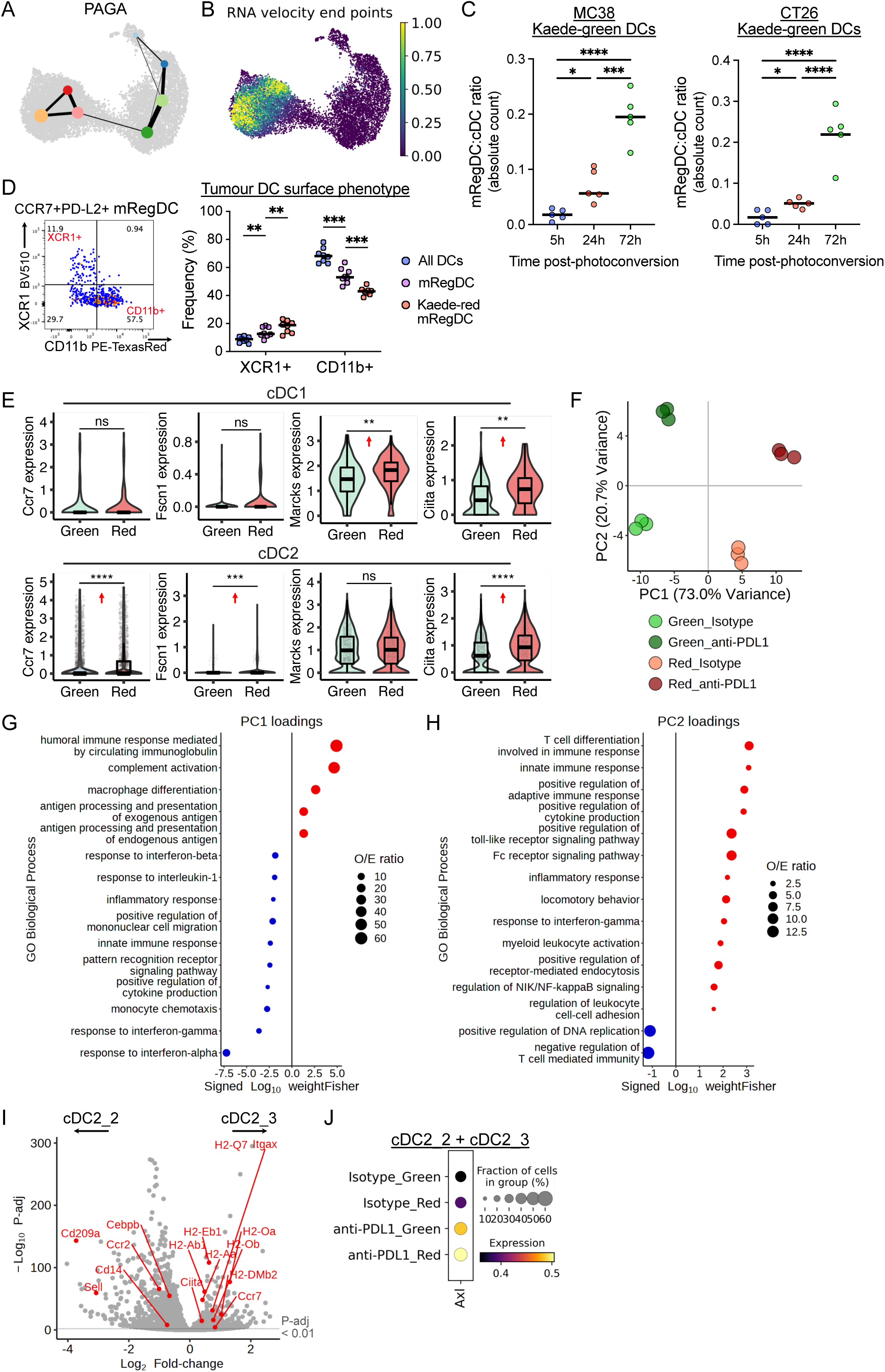
Maturation trajectory in tumour DCs. (A) Partition-based graph abstraction (PAGA) of scRNA-seq of tumour DCs. (B) Terminal states (end points) of RNA velocity analysis. (C) Flow cytometry of mRegDC to cDC ratio in Kaede-green DCs; time course post- photoconversion. MC38 and CT26 tumours were used; points represent independent mice; one-way ANOVA with Šidák’s multiple comparisons test was used. (D) Flow cytometry of surface cDC1 (XCR1^+^) and cDC2 (CD11b^+^) marker expression on mRegDCs from MC38-Ova tumours. Paired t-test with FDR correction was used. (E) Expression of selected DC migration genes and *Ciita* in scRNA-seq data of cDC1 and cDC2. Arrows indicate relative expression in Kaede-red versus Kaede-green cells; Wilcoxon rank-sum test was used. (F) Principal component (PC) analysis of scRNA-seq of cDC2s, pseudo-bulked to 3 artificial replicates per condition. PC1; variance due to Kaede profile. PC2; variance due to treatment. (G-H) Gene ontology (GO) term enrichment test for the top 100 loading genes of PC1 (G) and PC2 (H). O/E, observed/expected ratio; colour, direction of gene loading. Fisher-exact test was used. (I) Differential gene expression between cDC2_2 and cDC2_3. Selected DEGs relating to DC maturation, antigen presentation, MHC-II and migration are highlighted in red (all *P*-adj < 0.01). (J) Expression of *Axl* in cDC2_2 and cDC2_3 combined.

**Extended Data Figure 4.**
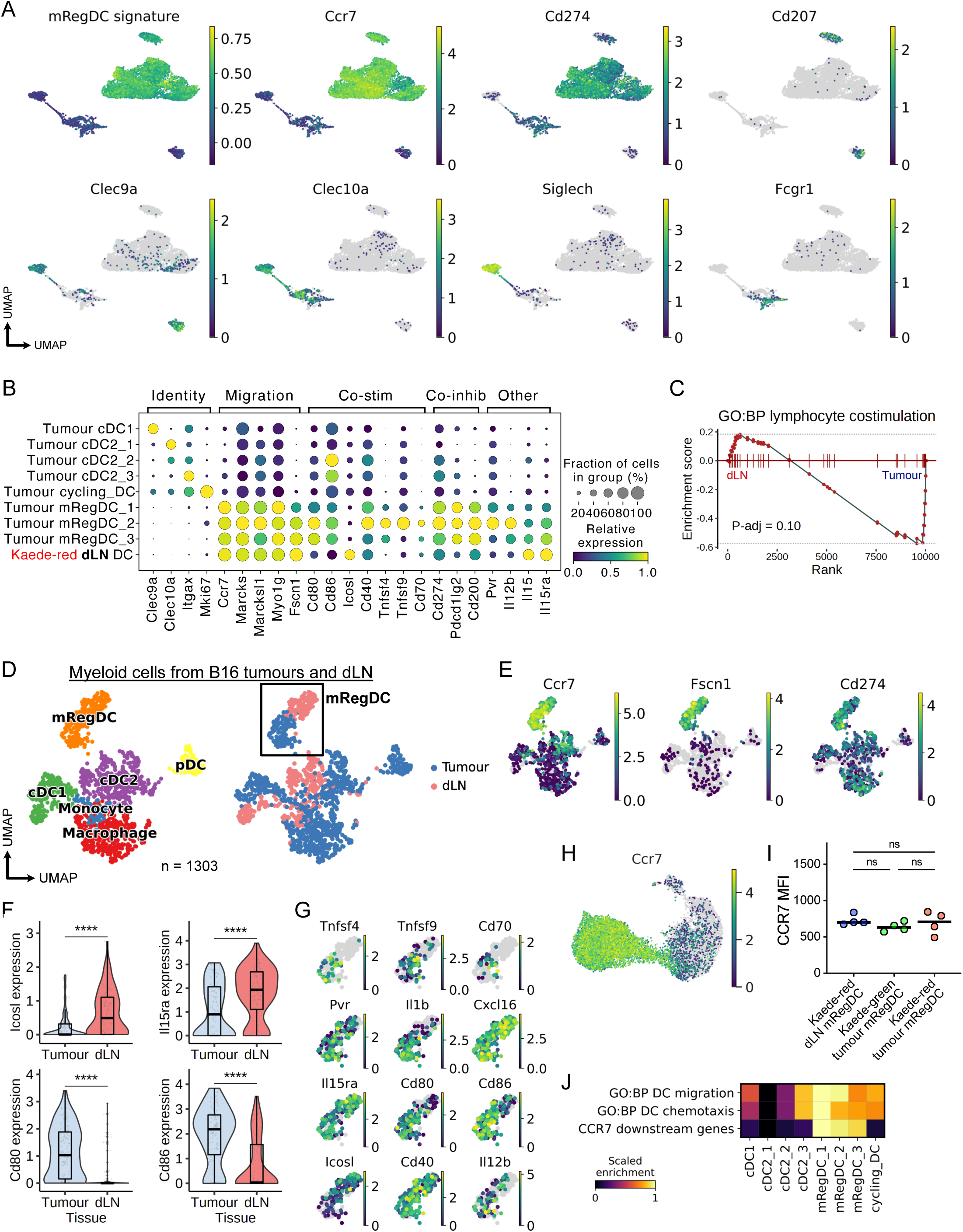
Comparison of tumour-residing mRegDCs versus LN mRegDC emigrants. (A) Expression of mRegDC or myeloid cell canonical marker genes in scRNA-seq of LN myeloid cells (Fig. 2D). (B) Expression of selected genes in tumour DCs (isotype control-treated) and Kaede-red DC emigrants in tumour-dLNs. (C) GSEA of “*GO:biological process (BP) lymphocyte co-stimulation (GO:0031294)*”, comparing Kaede-red mRegDCs from the dLN versus tumour mRegDCs. (D) UMAP of myeloid cells from scRNA-seq of murine subcutaneous B16 tumours and tumour-dLNs, by cell-type and tissue. (E) Expression of mRegDC genes in (D). (F) Violin plots of selected genes in scRNAseq of mRegDCs from B16 tumours and tumour- dLNs. Wilcoxon rank-sum test was used. (G) Expression of selected DEGs between in tumour and dLN mRegDCs from (D), indicated within the box. (H-I) Expression of CCR7 in DCs from MC38-Ova tumours in the scRNA-seq data (H) and by flow cytometry (I). Paired t-test with FDR correction was used (I). (J) Gene signature enrichment of pathways relating to DC movement and CCR7 signalling.

**Extended Data Figure 5.**
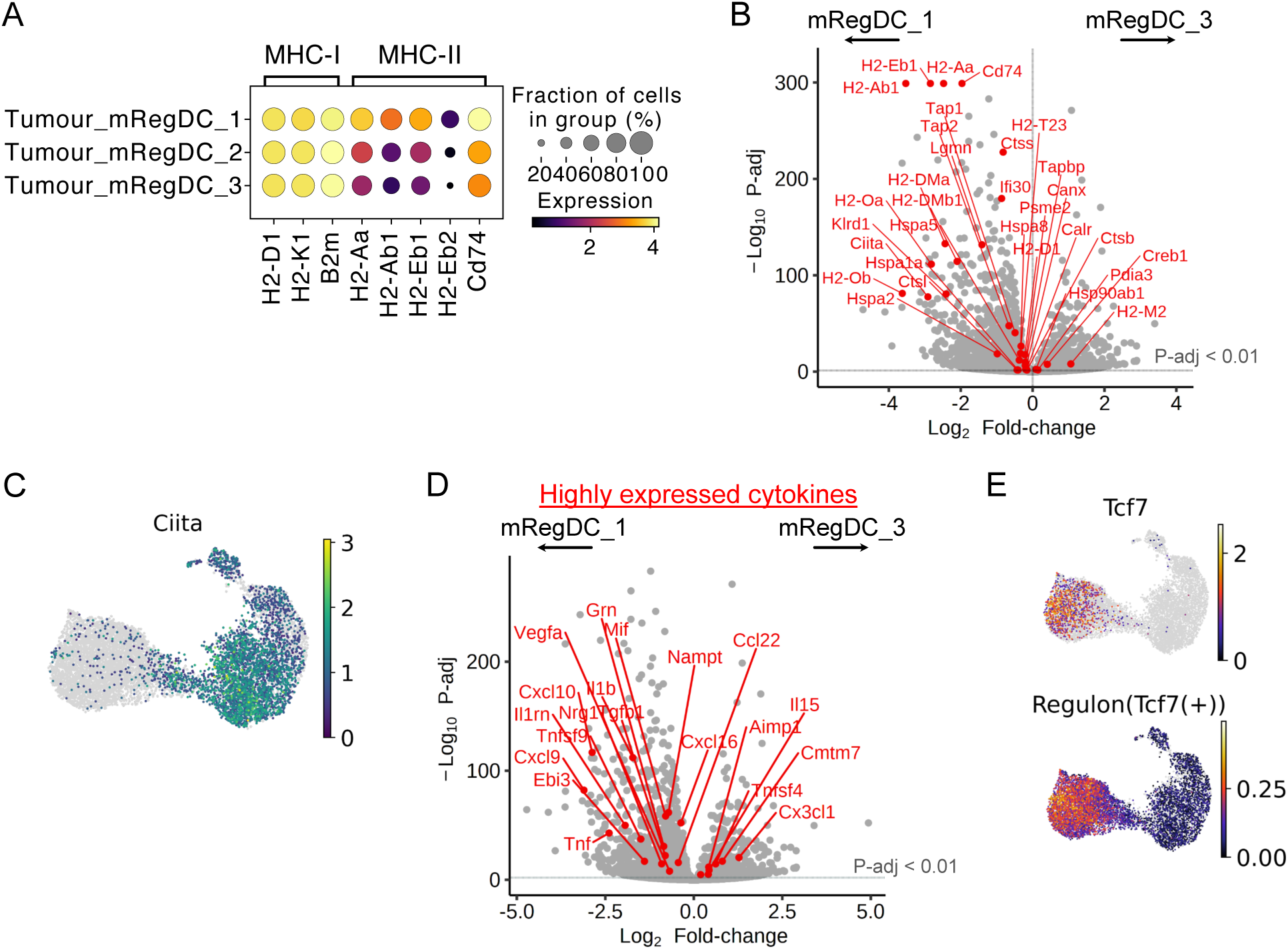
Transcriptional changes in mRegDCs with prolonged tumour residence. (A) Dot plot of class-I and class-II MHC gene expression (log-transformed, unscaled) in mRegDCs. (B) Differential gene expression between mRegDC_1 and mRegDC_3. Significant DEGs from “*KEGG antigen processing and presentation*” are highlighted in red (*P*-adj < 0.01). (C) Expression of *Ciita*. (D) Differential gene expression between mRegDC_1 and mRegDC_3. Significant DEGs from “*GO: molecular function (MF) cytokine activity (GO:0005125)*” are highlighted in red (*P*-adj < 0.01). (E) *Tcf7* expression and regulon activity score.

**Extended Data Figure 6.**
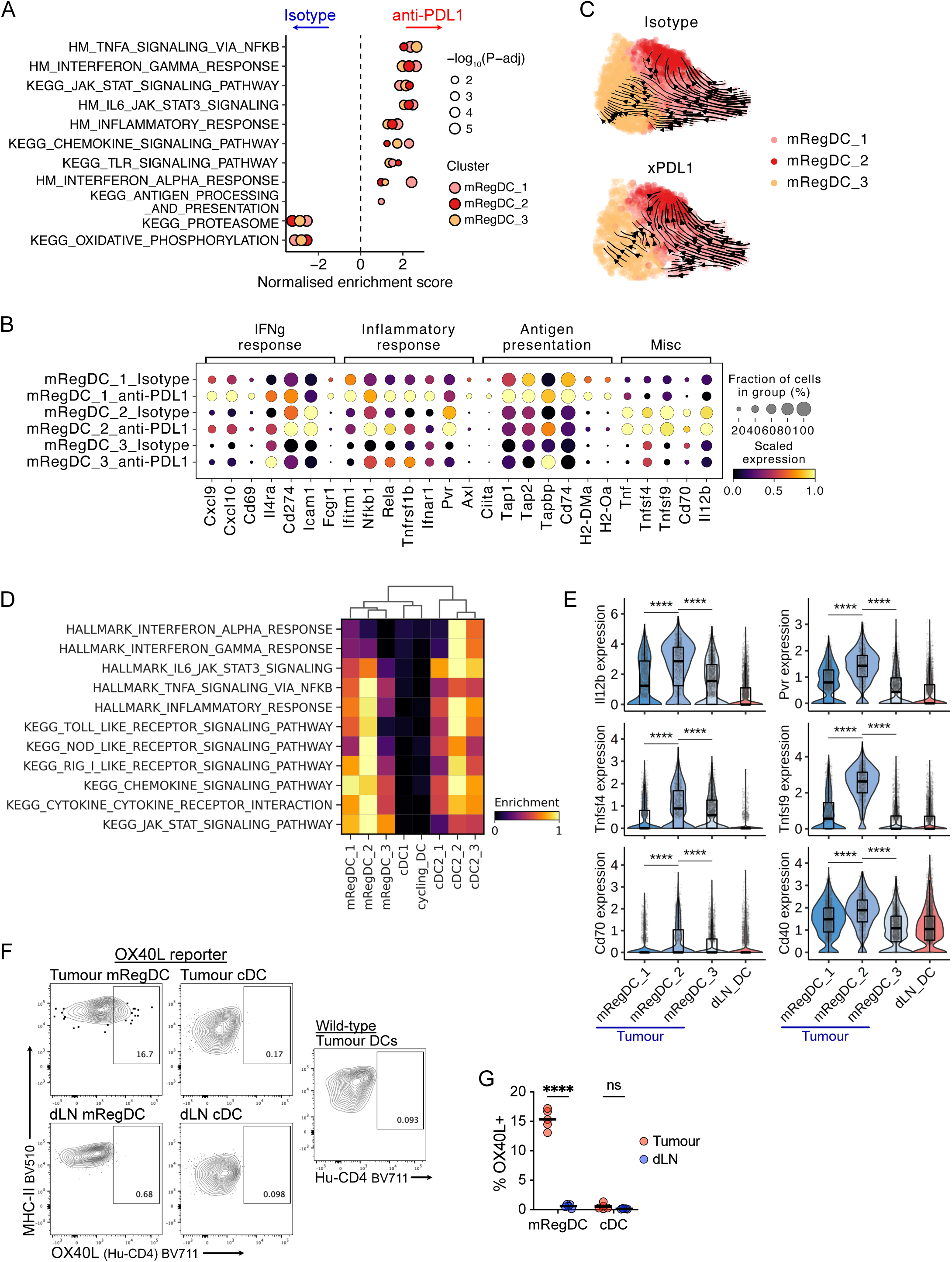
mRegDC activation following anti-PD-L1 treatment. (A) GSEA of mRegDCs in anti-PD-L1-treated versus isotype control-treated tumours; Hallmark (HM) and KEGG pathways. Only significant pathways (*P*-adj < 0.05) are shown. (B) Dot plot of selected leading-edge genes from GSEA analysis of “*Hallmark interferon gamma response*”, “*Hallmark inflammatory response*”, and “*KEGG antigen processing and presentation*” pathways from (A). (C) RNA velocity trajectory in tumour mRegDCs split by treatment group. (D) Gene signature scores of selected pathways in mRegDCs. (E) Expression of selected ligands differentially expressed between tumour mRegDCs and mRegDC emigrants in the dLN (Kaede-red). Wilcoxon rank-sum test was used. (F) Representative flow cytometry of surface OX40L expression (using OX40L^+/Human-CD4^ reporter mice) on DCs from MC38-Ova tumours and dLNs. Wild-type DC (non-reporter) were used for OX40L^+^ gating. (G) Quantification of (F). Points represent independent mice, paired t-test with FDR correction was used.

**Extended Data Figure 7.**
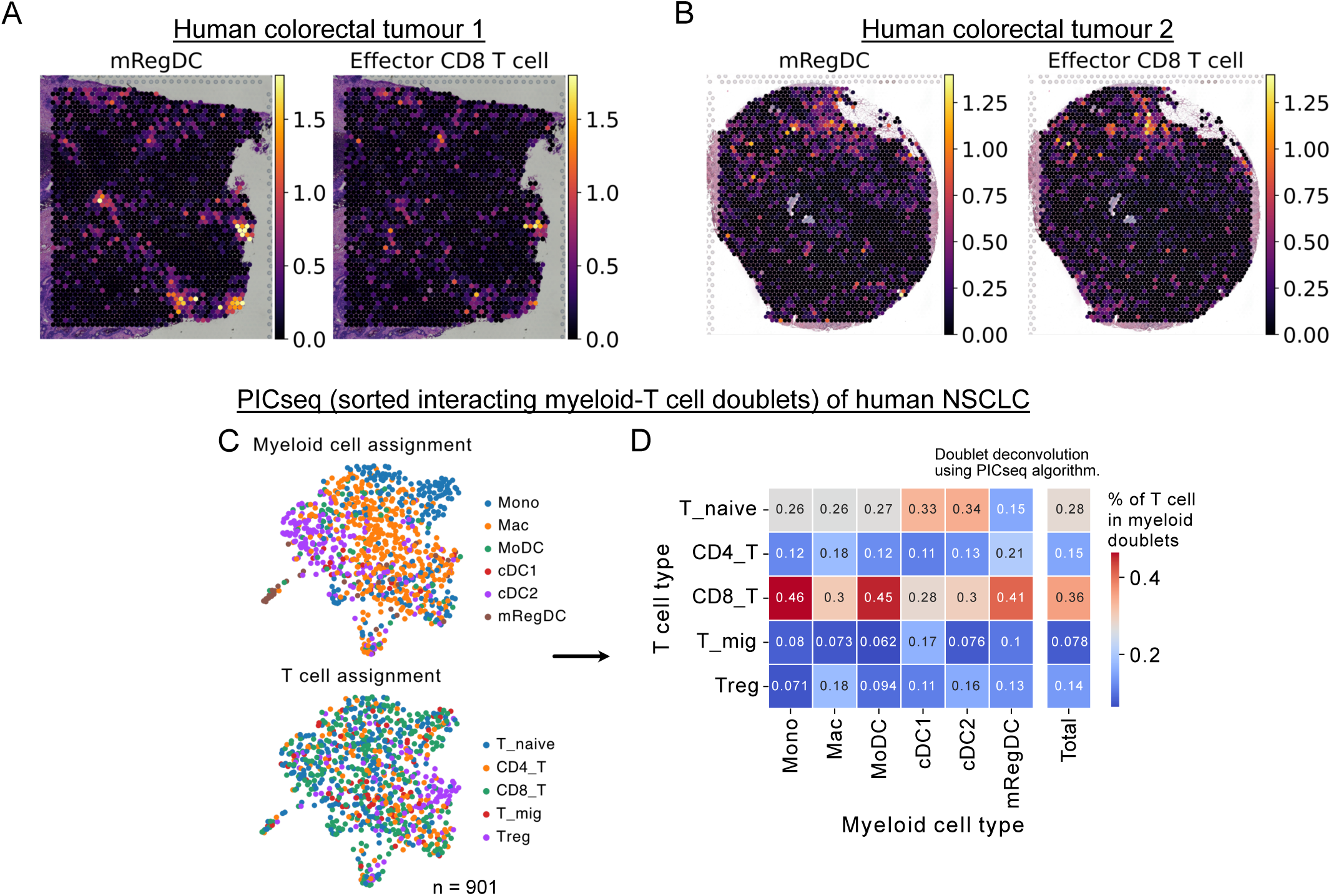
mRegDC-CD8^+^ T cell engagement in human solid tumours. (A-B) Gene signature scores for mRegDCs and effector CD8^+^ T cells in spatial transcriptomics (10X Genomics Visium) of independent human CRC tumour sections associated with Fig. 4D (*n=* 2). (C) Sequencing of physically interacting cells (PICseq, myeloid-T cell doublets) from human NSCLC (*n* = 10 patients). UMAP of PICs coloured by the deconvolved myeloid and T cell assignment for each doublet (PICseq algorithm). Each dot represents one myeloid-T cell doublet. (D) Heatmap of frequency (values) of myeloid-T cell doublet combinations in PICseq of NSCLC.

**Extended Data Figure 8.**
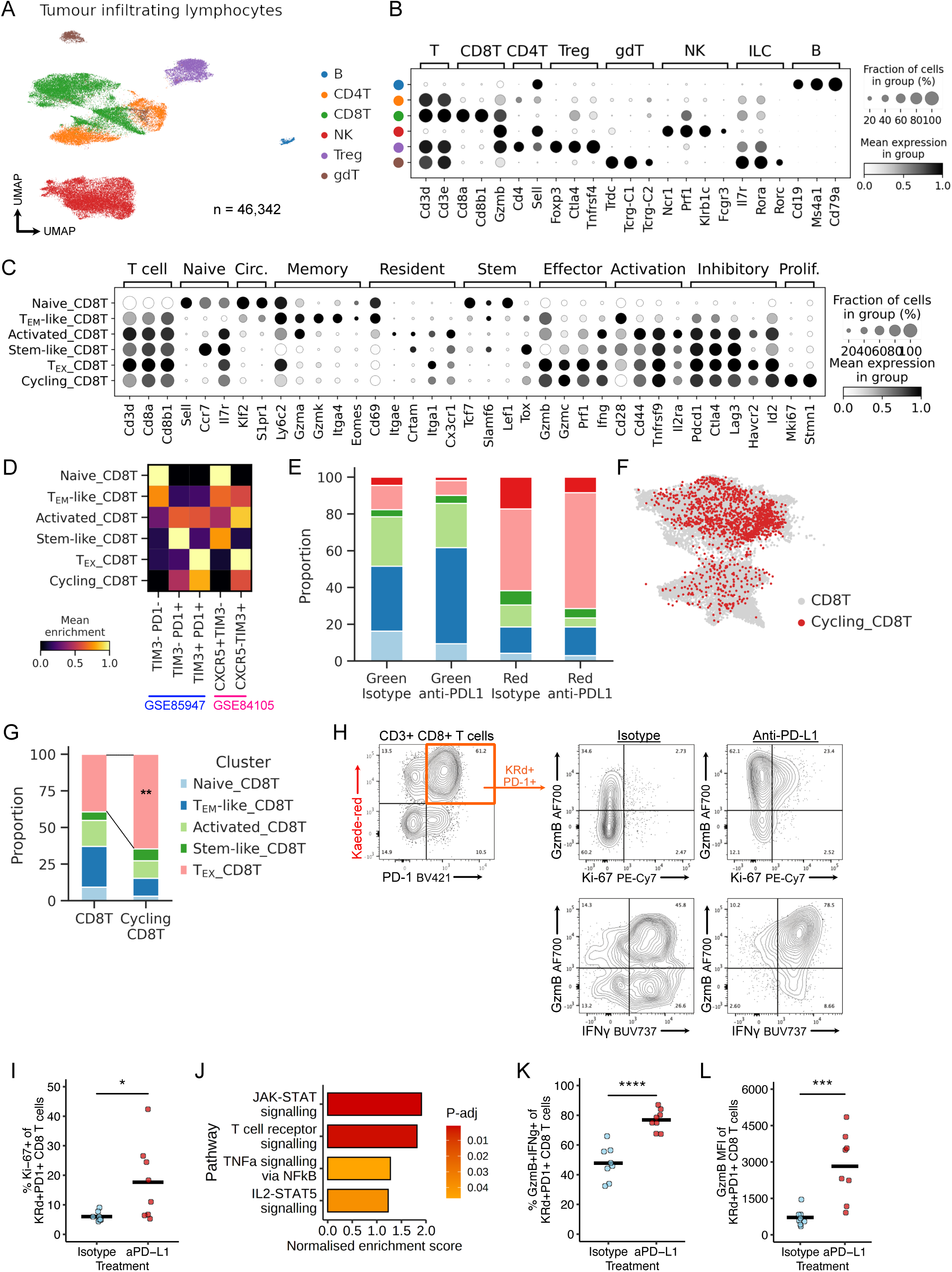
Single cell profiling of tumour CD8^+^ T cells following anti-PD- L1 treatment. (A) UMAP of TILs from scRNA-seq of FACS-sorted CD45^+^TER119^-^ Kaede-green^+^/Kaede- red^+^ cells 48h after photoconversion of subcutaneous MC38-Ova tumours, and canonical marker gene expression (B). (C) Canonical marker gene expression of CD8^+^ T cell clusters from Fig. 5A. (D) Gene signature enrichment of published T cell exhaustion transcriptomic signatures in CD8^+^ T cell clusters. (E) Proportion of CD8^+^ T cells by Kaede fluorescence and treatment groups. (F) Regression of cell cycle genes from the cycling_CD8T cluster followed by re-integration, and label transfer. Majority of the cycling cluster embedded with the CD8^+^ TEX cell cluster. (G) Proportion of cycling CD8^+^ T cells belonging to each CD8^+^ T cell cluster, from (F). Chi- squared test for over-representation of TEX CD8^+^ T cells was used. (H) Representative flow cytometry of Ki-67^+^, GzmB^+^ and IFNγ^+^ expression in CD3^+^CD8^+^Kaede-red^+^PD-1^+^ TEX cells, following anti-PD-L1 treatment. (I) Flow cytometry of Kaede-red^+^PD-1^+^CD8^+^ T cells, showing increased frequency of cycling cells (Ki-67^+^) following anti-PD-L1. (J) GSEA for TEX CD8^+^ T cells from anti-PD-L1 versus isotype control-treated tumours. (K) Flow cytometry of Kaede-red^+^PD-1^+^CD8^+^ T cells, showing increased frequency of GzmB^+^IFΝγ^+^ cells following anti-PD-L1. (L) Flow cytometry of Kaede-red^+^PD-1^+^ CD8^+^ T cells, showing increased expression (MFI) of GzmB following anti-PD-L1. Points represent independent mice, student’s t-test (I,K) or Mann-Whitney U test was used (L).

**Extended Data Figure 9.**
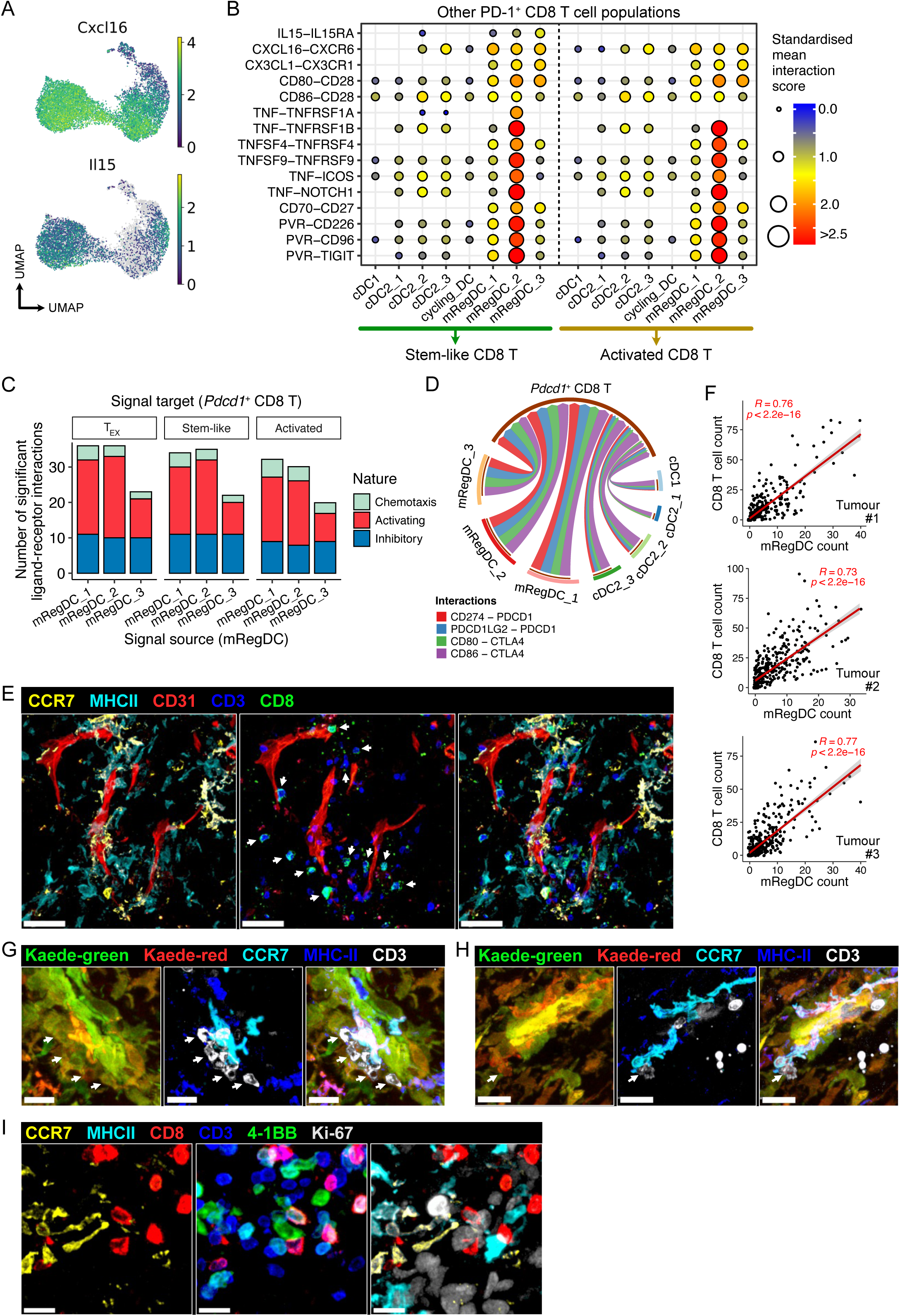
mRegDCs communicate with CD8^+^ T cells in murine tumours. (A) Expression of *Cxcl16* and *Il15*. (B-D) CellPhoneDB cell-cell communication analysis between DCs and *Pdcd1*^+^ CD8^+^ T cells in scRNA-seq of MC38-Ova tumours. (B) Ligand-receptor predicted interactions between tumour DCs and activated or stem-like CD8^+^cells. (C) Interactions that are well-described in existing literature to influence CD8^+^ T cells function were identified and classified based on whether engagement of the cognate T cell receptor was activating (increase in effector function, proliferation, or survival) or inhibitory in nature. (D) *PDCD1* or *CTLA4*-mediated inhibitory signals; edge width scaled to standardised interaction scores. Only significant interactions (*p* < 0.05) shown (B,D). (E) Representative confocal microscopy images of MC38 tumours, showing co-localisation of CCR7^+^MHC-II^+^ mRegDCs and CD3^+^CD8^+^ T cells (arrows). Scale bar, 80 μm. (F) Quantification of (E); spatial correlation (Pearson) between mRegDC and CD8^+^ T cell counts in 200 x 200 μm fields of MC38 tumour sections. 3 independent tumours were analysed. (G-H) Representative microscopy of independent MC38 tumours 48h after tumour photoconversion, showing interaction between Kaede-red CCR7^+^MHC-II^+^ mRegDCs and Kaede-red CD3^+^ T cells (arrows). Scale bar, 15 μm (G), 20 μm (H). (I) Representative microscopy of independent MC38 tumour, showing co-localisation of CCR7^+^MHC-II^+^ mRegDCs and CD3^+^CD8^+^4-1BB^+^Ki-67^+^ T cells. Scale bar, 15 μm.

**Supplementary Figure 10.**
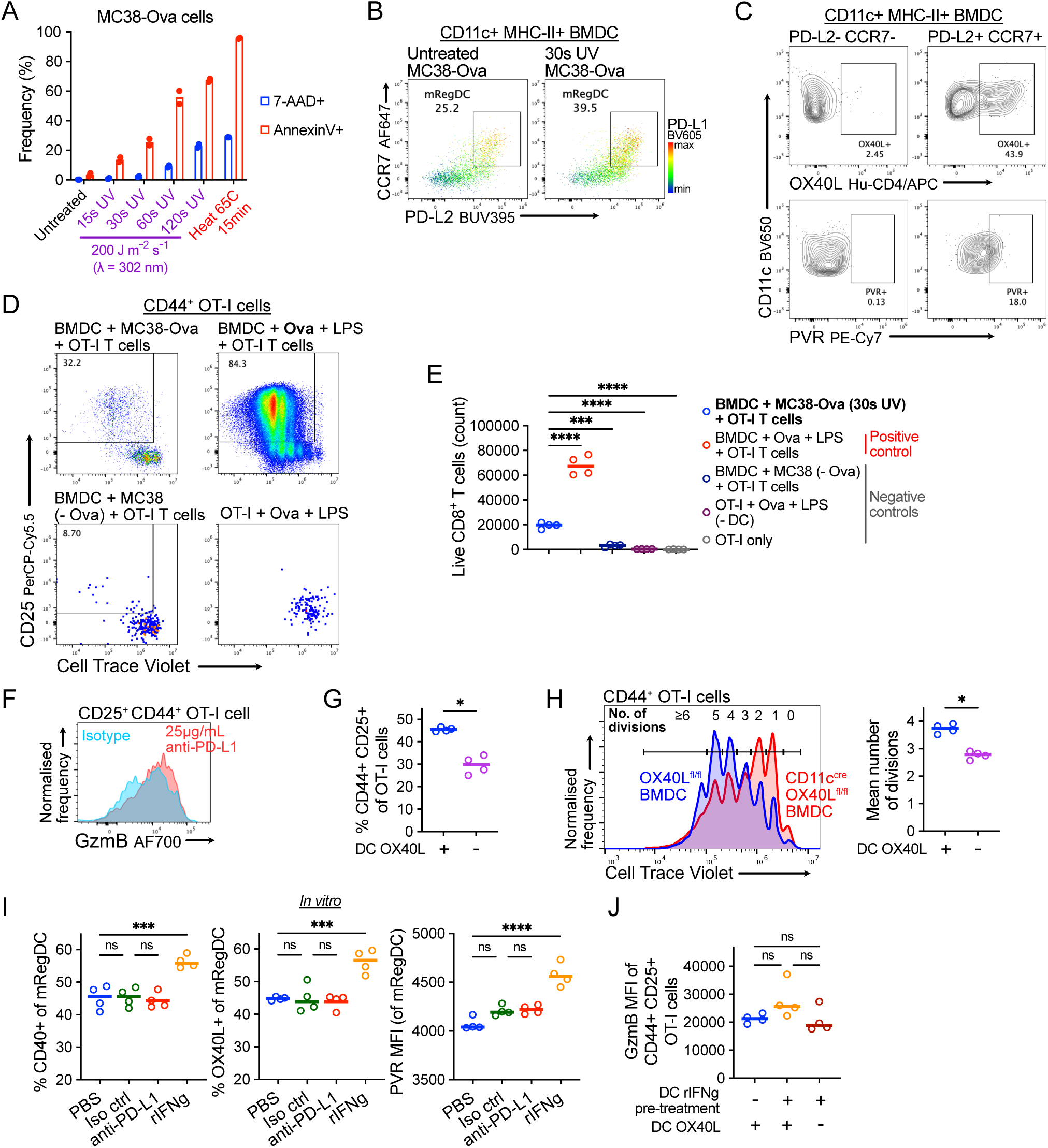
*In vitro* DC OT-I CD8^+^ T cell co-culture. (A) Cell death following UV irradiation of MC38-Ova cell monolayer *in vitro*; for optimisation of tumour cell-line apoptosis. 30s UV exposure was chosen for subsequent experiments. (B) Representative flow cytometry of BMDC following 8h culture with UV-irradiated MC38-Ova cells. (C) Representative flow cytometry OX40L and PVR expression on BMDC. (D) Representative flow cytometry of OT-I activation and proliferation in DC co-cultures. Culture with apoptotic MC38-Ova experienced FACS-sorted PD-L2^+^CCR7^+^ BMDC (mReg-BMDC), top left; positive control (BMDC + Ova + LPS), top right; negative controls, bottom left (no Ova antigen) and right (no DC). (E) Number of live CD8^+^ T cells in mReg-BMDC + OT-I co-culture set-up versus controls. (F) Representative flow cytometry of GzmB expression in OT-I cells; +/- anti-PD-L1 antibodies. (G) Flow cytometry of OT-I activation, and (H) CTV proliferation in OT-I cells and mean number of cell divisions, following culture with OX40L-expressing (+) or OX40L-deficient (-) mReg-BMDC. (I) Flow cytometry of mReg-BMDCs following 8h treatment with antibodies or recombinant IFNγ (before OT-I co-culture). (J) Flow cytometry of OT-I cells following co-culture with mReg-BMDC; DCs pre-treated with recombinant IFNγ (+) or PBS (-), OX40L-expressing (+) or OX40L-deficient (-) BMDCs. One-way ANOVA with Šidák’s multiple comparisons test was used (E-J).

**Extended Data Figure 11.**
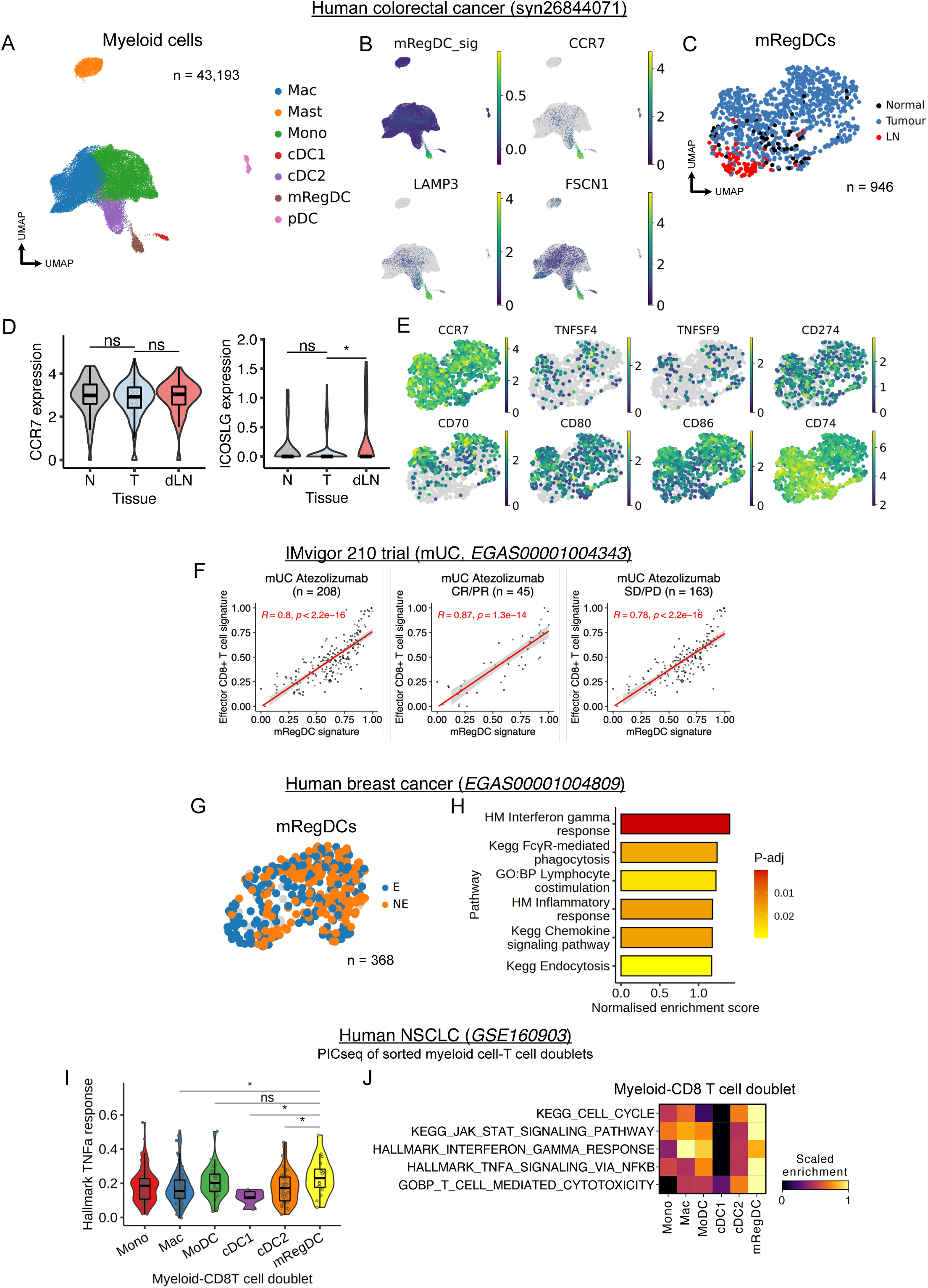
mRegDC heterogeneity and interaction with CD8^+^ T cells in human cancer. (A) UMAP of myeloid cells from scRNA-seq of human CRC (*n* = 63 patients) with paired tumour (T), normal adjacent tissue (N) and dLN samples, and expression of mRegDC signature genes (B). (C) UMAP of mRegDCs subset from (A), coloured by tissue origin. (D) Expression of *CCR7* and *ICOSLG* in mRegDCs by tissue. Wilcoxon rank-sum test was used. (E) Expression of selected genes in mRegDCs from (C). Molecules associated with tumour- residing mRegDCs in mice were also preferentially expressed in tumour mRegDCs in human CRC, but not the dLN. (F) Pearson correlation between mRegDC signature genes and effector CD8^+^ T cell signature genes in bulk RNA-seq of 208 mUC tumours treated with atezolizumab (IMvigor 210 trial). Left, middle and right panel show all patients, clinical responders, and non-responders respectively. (G) UMAP of mRegDCs from human breast cancer (*n =* 29 patients). (H) GSEA of mRegDCs from T cell clonotype expanders (E, i.e. responders, *n* = 9 patients) versus non-expanders (NE, i.e. non-responders, *n* = 20 patients), in breast tumours treated with anti-PD-1. (I) Gene signature score for “*Hallmark TNFa response via NFkB*” in PICseq of NSCLC, grouped by the myeloid cell identity in each myeloid-T cell doublet. Wilcoxon rank-sum test was used. (J) Gene signature scores of selected pathways in PICs, grouped by the myeloid cell identity.

**Extended Data Figure 12.**
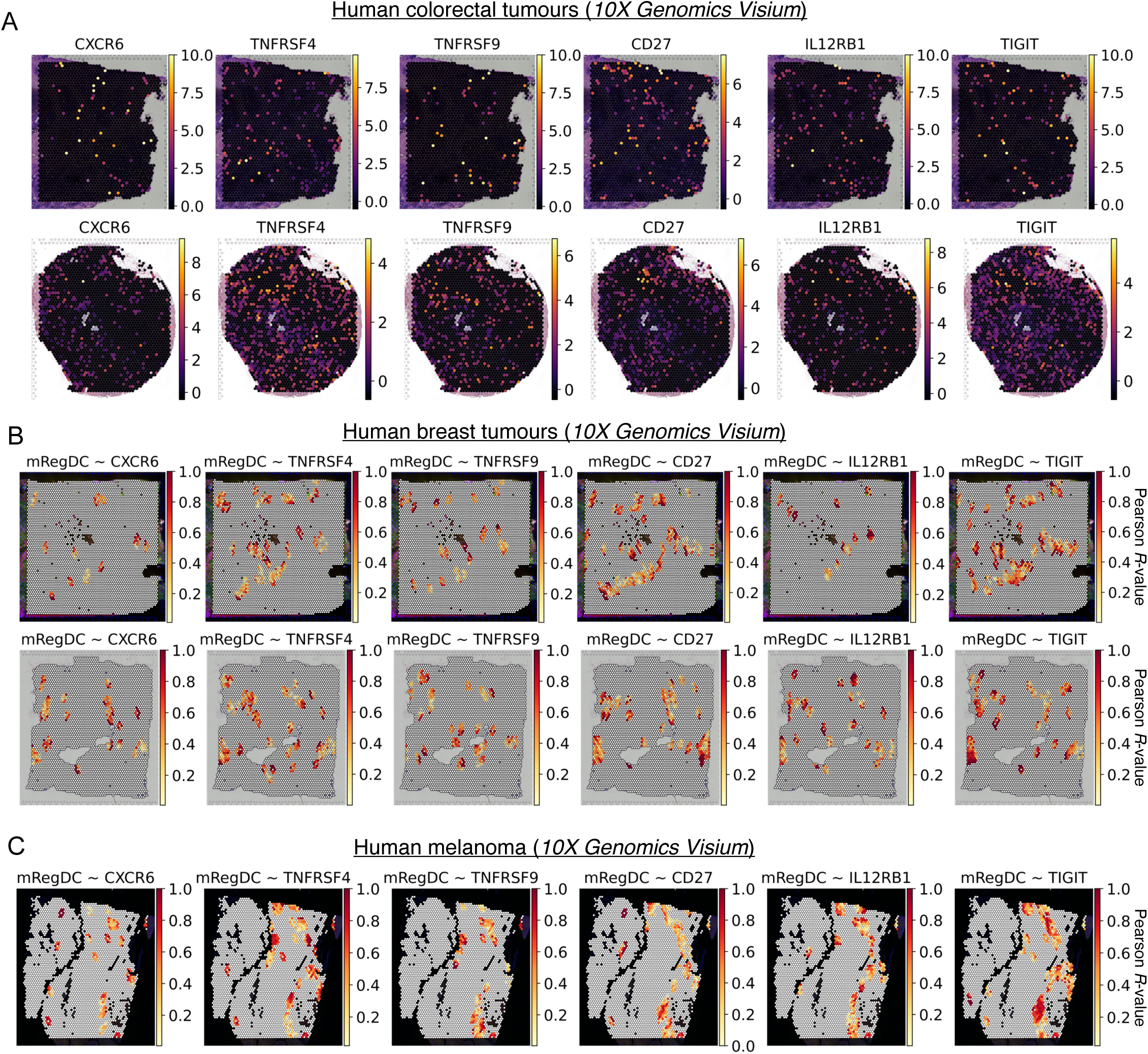
Visium spatial transcriptomics of human cancer. (A) Expression of selected receptors expressed by CD8^+^ T cells which facilitate interactions with tumour- residing mRegDCs in spatial transcriptomics (10X Genomics Visium) of independent human CRC tumour sections (*n= 2*). (B-C) Spatial correlation (Pearson R-value) of mRegDC signature scores and selected mRegDC-ligand receptors expressed by CD8^+^ T cells, in spatial transcriptomics of independent human breast tumour sections (B, *n* = 2) and a human melanoma section (C, *n* = 1).

## References

1. Wculek, S. K. et al. Dendritic cells in cancer immunology and immunotherapy. Nature Reviews Immunology 2019 20:1 20, 7–24 (2019).

2. Roberts, E. W. et al. Critical Role for CD103(+)/CD141(+) Dendritic Cells Bearing CCR7 for Tumor Antigen Trafficking and Priming of T Cell Immunity in Melanoma. Cancer Cell 30, 324–336 (2016).

3. Sallusto, F. et al. Rapid and coordinated switch in chemokine receptor expression during dendritic cell maturation. Eur J Immunol 28, 2760–2769 (1998).

4. Böttcher, J. P. & Reis e Sousa, C. The Role of Type 1 Conventional Dendritic Cells in Cancer Immunity. Trends Cancer 4, 784–792 (2018).

5. Maier, B. et al. A conserved dendritic-cell regulatory program limits antitumour immunity. Nature 580, 257–262 (2020).

6. 6. Kvedaraite, E. & Ginhoux, F. Human dendritic cells in cancer. Sci Immunol 7, (2022).

7. Ardouin, L. et al. Broad and Largely Concordant Molecular Changes Characterize Tolerogenic and Immunogenic Dendritic Cell Maturation in Thymus and Periphery. Immunity 45, 305–318 (2016).

8. Ginhoux, F., Guilliams, M. & Merad, M. Expanding dendritic cell nomenclature in the single- cell era. Nature Reviews Immunology 2022 22:2 22, 67–68 (2022).

9. Gerhard, G. M., Bill, R., Messemaker, M., Klein, A. M. & Pittet, M. J. Tumor-infiltrating dendritic cell states are conserved across solid human cancers. Journal of Experimental Medicine 218, (2021).

10. Cheng, S. et al. A pan-cancer single-cell transcriptional atlas of tumor infiltrating myeloid cells. Cell 184, 792–809.e23 (2021).

11. Zilionis, R. et al. Single-Cell Transcriptomics of Human and Mouse Lung Cancers Reveals Conserved Myeloid Populations across Individuals and Species. Immunity 50, 1317–1334.e10 (2019).

12. Goc, J. et al. Dendritic cells in tumor-associated tertiary lymphoid structures signal a th1 cytotoxic immune contexture and license the positive prognostic value of infiltrating CD8+ t cells. Cancer Res 74, 705–715 (2014).

13. Movassagh, M. et al. Selective Accumulation of Mature DC-Lamp+ Dendritic Cells in Tumor Sites Is Associated with Efficient T-Cell-Mediated Antitumor Response and Control of Metastatic Dissemination in Melanoma. Cancer Res 64, 2192–2198 (2004).

14. de Mingo Pulido, Á., et al. TIM-3 Regulates CD103+ Dendritic Cell Function and Response to Chemotherapy in Breast Cancer. Cancer Cell 33, 60–74.e6 (2018).

15. Dieu-Nosjean, M. C. et al. Long-term survival for patients with non-small-cell lung cancer with intratumoral lymphoid structures. J Clin Oncol 26, 4410–4417 (2008).

16. Peng, Q. et al. PD-L1 on dendritic cells attenuates T cell activation and regulates response to immune checkpoint blockade. Nature Communications 2020 11:1 11, 1–8 (2020).

17. Oh, S. A. et al. PD-L1 expression by dendritic cells is a key regulator of T-cell immunity in cancer. Nature Cancer 2020 1:7 1, 681–691 (2020).

18. Zhang, Q. et al. Landscape and Dynamics of Single Immune Cells in Hepatocellular Carcinoma. Cell 179, 829–845.e20 (2019).

19. Li, Z. et al. In vivo labeling reveals continuous trafficking of TCF-1+ T cells between tumor and lymphoid tissue. J Exp Med 219, (2022).

20. Wherry, E. J. T cell exhaustion. Nat Immunol 12, 492–499 (2011).

21. Platonova, S. et al. Profound coordinated alterations of intratumoral NK cell phenotype and function in lung carcinoma. Cancer Res 71, 5412–5422 (2011).

22. 22. Dean, I. et al. Rapid establishment of a tumor-retained state curtails the contribution of conventional NK cells to anti-tumor immunity in solid cancers. bioRxiv 2023.08.10.552797 (2023) doi:10.1101/2023.08.10.552797.

23. 23. The Cancer Genoma Atlas. TCGA. National Cancer Institute (NCI) and National Human Genome Research Institute (NHGRI) (2013).

24. Bassez, A. et al. A single-cell map of intratumoral changes during anti-PD1 treatment of patients with breast cancer. Nat Med 27, 820–832 (2021).

25. Li, H. et al. Dysfunctional CD8 T Cells Form a Proliferative, Dynamically Regulated Compartment within Human Melanoma. Cell 176, 775–789.e18 (2019).

26. Pelka, K. et al. Spatially organized multicellular immune hubs in human colorectal cancer. Cell 184, 4734–4752.e20 (2021).

27. Curtis, C. et al. The genomic and transcriptomic architecture of 2,000 breast tumours reveals novel subgroups. Nature 486, 346–352 (2012).

28. Tomura, M. et al. Monitoring cellular movement in vivo with photoconvertible fluorescence protein ‘Kaede’ transgenic mice. Proc Natl Acad Sci U S A 105, 10871–10876 (2008).

29. Riley, J. L., Westerheide, S. D., Price, J. A., Brown, J. A. & Boss, J. M. Activation of class II MHC genes requires both the X ☐ region and the class II transactivator (CIITA). Immunity 2, 533–543 (1995).

30. Miller, J. C. et al. Deciphering the transcriptional network of the DC lineage. Nat Immunol 13, 888 (2012).

31. Riley, J. L. et al. Modulation of TCR-induced transcriptional profiles by ligation of CD28, ICOS, and CTLA-4 receptors. Proc Natl Acad Sci U S A 99, 11790–11795 (2002).

32. Garris, C. S. et al. Successful Anti-PD-1 Cancer Immunotherapy Requires T Cell-Dendritic Cell Crosstalk Involving the Cytokines IFN-γ and IL-12. Immunity 49, 1148–1161.e7 (2018).

33. Dubois, S. P., Waldmann, T. A. & Müller, J. R. Survival adjustment of mature dendritic cells by IL-15. Proc Natl Acad Sci U S A 102, 8662 (2005).

34. Davidson, S. et al. Single-Cell RNA Sequencing Reveals a Dynamic Stromal Niche That Supports Tumor Growth. Cell Rep 31, (2020).

35. Croft, M. The role of TNF superfamily members in T-cell function and diseases. Nature Reviews Immunology 2009 9:4 9, 271–285 (2009).

36. Johnston, R. J. et al. The Immunoreceptor TIGIT Regulates Antitumor and Antiviral CD8+ T Cell Effector Function. Cancer Cell 26, 923–937 (2014).

37. 10x Genomics. 10x Genomics Spatial Gene Expression. Visium demonstration dataset https://www.10xgenomics.com/resources/datasets (2022).

38. Cohen, M. et al. The interaction of CD4+ helper T cells with dendritic cells shapes the tumor microenvironment and immune checkpoint blockade response. Nat Cancer 3, 303–317 (2022).

39. Im, S. J. et al. Defining CD8+ T cells that provide the proliferative burst after PD-1 therapy. Nature 537, 417–421 (2016).

40. Singer, M. et al. A Distinct Gene Module for Dysfunction Uncoupled from Activation in Tumor- Infiltrating T Cells. Cell 166, 1500–1511.e9 (2016).

41. di Pilato, M. et al. CXCR6 positions cytotoxic T cells to receive critical survival signals in the tumor microenvironment. Cell 184, 4512–4530.e22 (2021).

42. Joanito, I. et al. Single-cell and bulk transcriptome sequencing identifies two epithelial tumor cell states and refines the consensus molecular classification of colorectal cancer. Nature Genetics 2022 54:7 54, 963–975 (2022).

43. Rosenberg, J. E. et al. Atezolizumab in patients with locally advanced and metastatic urothelial carcinoma who have progressed following treatment with platinum-based chemotherapy: a single-arm, multicentre, phase 2 trial. Lancet 387, 1909–1920 (2016).

44. Banchereau, R. et al. Molecular determinants of response to PD-L1 blockade across tumor types. Nature Communications 2021 12:1 12, 1–11 (2021).

45. Balar, A. v., et al. Atezolizumab as first-line treatment in cisplatin-ineligible patients with locally advanced and metastatic urothelial carcinoma: a single-arm, multicentre, phase 2 trial. The Lancet 389, 67–76 (2017).

46. Chen, Z. et al. Single-cell RNA sequencing highlights the role of inflammatory cancer- associated fibroblasts in bladder urothelial carcinoma. Nature Communications 2020 11:1 11, 1–12 (2020).

47. Sun, D. et al. Identifying phenotype-associated subpopulations by integrating bulk and single- cell sequencing data. Nature Biotechnology 2021 40:4 40, 527–538 (2021).

48. Prokhnevska, N. et al. CD8+ T cell activation in cancer comprises an initial activation phase in lymph nodes followed by effector differentiation within the tumor. Immunity 56, 107–124.e5 (2022).

49. Thumkeo, D. et al. PGE2-EP2/EP4 signaling elicits immunosuppression by driving the mregDC-Treg axis in inflammatory tumor microenvironment. Cell Rep 39, (2022).

50. Magen, A. et al. Intratumoral dendritic cell–CD4+ T helper cell niches enable CD8+ T cell differentiation following PD-1 blockade in hepatocellular carcinoma. Nature Medicine 2023 29:6 29, 1389–1399 (2023).

51. Tang, F. et al. A pan-cancer single-cell panorama of human natural killer cells. Cell (2023) doi:10.1016/J.CELL.2023.07.034.

52. Yi, M. et al. Combination strategies with PD-1/PD-L1 blockade: current advances and future directions. Molecular Cancer 2022 21:1 21, 1–27 (2022).

53. Cho, B. C. et al. Tiragolumab plus atezolizumab versus placebo plus atezolizumab as a first-line treatment for PD-L1-selected non-small-cell lung cancer (CITYSCAPE): primary and follow- up analyses of a randomised, double-blind, phase 2 study. Lancet Oncol 23, 781–792 (2022).

54. Banta, K. L. et al. Mechanistic convergence of the TIGIT and PD-1 inhibitory pathways necessitates co-blockade to optimize anti-tumor CD8+ T cell responses. Immunity 55, 512–526.e9 (2022).

55. Lee, J. v., et al. Combinatorial immunotherapies overcome MYC-driven immune evasion in triple negative breast cancer. Nature Communications 2022 13:1 13, 1–12 (2022).

56. Pichler, A. C. et al. TCR-independent CD137 (4-1BB) signaling promotes CD8+-exhausted T cell proliferation and terminal differentiation. Immunity (2023) doi:10.1016/J.IMMUNI.2023.06.007.

57. Gato-Cañas, M. et al. PDL1 Signals through Conserved Sequence Motifs to Overcome Interferon-Mediated Cytotoxicity. Cell Rep 20, 1818–1829 (2017).

58. Lucas, E. D. et al. PD-L1 Reverse Signaling in Dermal Dendritic Cells Promotes Dendritic Cell Migration Required for Skin Immunity. Cell Rep 33, (2020).

59. Whyte, C. E. et al. ACKR4 restrains antitumor immunity by regulating CCL21. Journal of Experimental Medicine 217, (2020).

60. Cheng, H. W. et al. CCL19-producing fibroblastic stromal cells restrain lung carcinoma growth by promoting local antitumor T-cell responses. Journal of Allergy and Clinical Immunology 142, 1257–1271.e4 (2018).

61. Dixon, K. O. et al. TIM-3 restrains anti-tumour immunity by regulating inflammasome activation. Nature 595, 101 (2021).

62. Radtke, A. J. et al. IBEX: A versatile multiplex optical imaging approach for deep phenotyping and spatial analysis of cells in complex tissues. Proc Natl Acad Sci U S A 117, 33455–33465 (2020).

63. Bankhead, P. et al. QuPath: Open source software for digital pathology image analysis. Scientific Reports 2017 7:1 7, 1–7 (2017).

64. Wolf, F. A., Angerer, P. & Theis, F. J. SCANPY: Large-scale single-cell gene expression data analysis. Genome Biol 19, 1–5 (2018).

65. Wolock, S. L., Lopez, R. & Klein, A. M. Scrublet: Computational Identification of Cell Doublets in Single-Cell Transcriptomic Data. Cell Syst 8, 281–291.e9 (2019).

66. Aibar, S. et al. SCENIC: single-cell regulatory network inference and clustering. Nature Methods 2017 14:11 14, 1083–1086 (2017).

67. Wolf, F. A. et al. PAGA: graph abstraction reconciles clustering with trajectory inference through a topology preserving map of single cells. Genome Biol 20, 1–9 (2019).

68. Setty, M. et al. Characterization of cell fate probabilities in single-cell data with Palantir. Nat Biotechnol 37, 451–460 (2019).

69. Bergen, V., Lange, M., Peidli, S., Wolf, F. A. & Theis, F. J. Generalizing RNA velocity to transient cell states through dynamical modeling. Nature Biotechnology 2020 38:12 38, 1408–1414 (2020).

70. Dann, E., Henderson, N. C., Teichmann, S. A., Morgan, M. D. & Marioni, J. C. Differential abundance testing on single-cell data using k-nearest neighbor graphs. Nat Biotechnol 40, 245– 253 (2022).

71. Efremova, M., Vento-Tormo, M., Teichmann, S. A. & Vento-Tormo, R. CellPhoneDB: inferring cell–cell communication from combined expression of multi-subunit ligand–receptor complexes. Nature Protocols 2020 15:4 15, 1484–1506 (2020).

72. Squair, J. W. et al. Confronting false discoveries in single-cell differential expression. Nature Communications 2021 12:1 12, 1–15 (2021).

73. Love, M. I., Huber, W. & Anders, S. Moderated estimation of fold change and dispersion for RNA-seq data with DESeq2. Genome Biol 15, (2014).

74. Subramanian, A. et al. Gene set enrichment analysis: A knowledge-based approach for interpreting genome-wide expression profiles. Proc Natl Acad Sci U S A 102, 15545–15550 (2005).

75. Alexa, A. & Rahnenfuhrer, J. topGO: Enrichment Analysis for Gene Ontology. Preprint at (2022).

76. Giladi, A. et al. Dissecting cellular crosstalk by sequencing physically interacting cells. Nature Biotechnology 2020 38:5 38, 629–637 (2020).

77. Aran, D., Hu, Z. & Butte, A. J. xCell: Digitally portraying the tissue cellular heterogeneity landscape. Genome Biol 18, 220 (2017).

78. Barbie, D. A. et al. Systematic RNA interference reveals that oncogenic KRAS-driven cancers require TBK1. Nature 462, 108–112 (2009).

79. Tuong, Z. K. et al. Resolving the immune landscape of human prostate at a single-cell level in health and cancer. Cell Rep 37, (2021).

